# Intramolecular autoinhibition of human PEX13 modulates peroxisomal import

**DOI:** 10.1101/2022.12.19.520972

**Authors:** Stefan Gaussmann, Julia Ott, Krzysztof M. Zak, Florent Delhommel, Grzegorz M. Popowicz, Wolfgang Schliebs, Ralf Erdmann, Michael Sattler

**Affiliations:** Bavarian NMR Center, Department of Bioscience, School of Natural Sciences, Technical University of Munich, Lichtenbergstr. 4, 85747 Garching, Germany; Institute of Structural Biology, Molecular Targets and Therapeutics Center, Helmholtz Zentrum München, Ingolstädter Landstr. 1, 85764 Neuherberg, Germany; Institute of Biochemistry and Pathobiochemistry, Department of Systems Biology, Faculty of Medicine, Ruhr University Bochum, 44780 Bochum, Germany

**Keywords:** peroxisome biogenesis, PEX13, peroxisomal matrix import, NMR spectroscopy, X-ray crystallography

## Abstract

Targeting and import of peroxisomal proteins depends on PEX5, PEX14 and PEX13. We present a biochemical and structural characterization of the PEX13 C-terminal region. By combining NMR spectroscopy, X-ray crystallography and biochemical methods, we show that the PEX13 SH3 domain mediates intramolecular interactions with a newly identified proximal FxxxF motif and also binds to WxxxF peptide motifs from the PEX5 NTD, demonstrating evolutionary conservation of this interaction from yeast to human. Strikingly, the C-terminal FxxxF motif autoinhibits the WxxxF/Y binding surface on the PEX13 SH3 domain. This is supported by high-resolution crystal structures, which show FxxxF or WxxxF/Y binding to the same, non-canonical surface on the SH3 domain. The FxxxF motif also binds the PEX14 NTD with micromolar affinity. Surprisingly, the canonical binding surface for PxxP motifs on the human PEX13 SH3 fold does not recognize PxxP motifs in PEX14, distinct from the yeast ortholog. The dynamic network of PEX13, PEX14 and PEX5 interactions mediated by diaromatic peptide motifs fine-tunes and modulates peroxisomal matrix import in cells.

## Introduction

Peroxisomes are single membrane enveloped organelles of eukaryotic cells which are essential for several metabolic pathways mainly related to lipid metabolism and to the removal of toxic oxidation products (Erdmann *et al*, 1997; Fujiki & Lazarow, 1985; Wanders, 2004; Wanders & Waterham, 2006). The physiological importance of these highly conserved organelles is emphasized by diseases such as Zellweger Spectrum Disorders (ZSD) that result from defects in peroxisome biogenesis (Klouwer *et al*, 2015). Biogenesis and function of peroxisomes rely on peroxisome-related proteins called peroxins (Distel *et al*, 1996; Ma *et al*, 2011) that are involved in membrane–assembly and post-translational matrix protein import into the organelle (Purdue & Lazarow, 2001). Human and yeast peroxins are abbreviated as “PEX” and “Pex”, respectively. Malfunction of PEX13, a peroxin crucial for peroxisomal import, leads to impaired biogenesis and neonatal death (Maxwell *et al*, 2003). The general mechanisms of peroxisomal biogenesis and matrix protein import are evolutionary conserved. Within the biogenesis process, cytosolic expressed peroxisomal membrane proteins of the translocation machinery are inserted into the peroxisomal membrane by the chaperone and receptor system PEX19-PEX3 (Hettema *et al*, 2000; Hoepfner *et al*, 2005; Sacksteder *et al*, 2000). Peroxisomal cargo proteins with their destination in the peroxisomal matrix possess conserved peroxisomal targeting signals, at their C-terminus (PTS1) or N-terminus (PTS2) (Ghosh & Berg, 2010; Gould *et al*, 1987). Cytosolic PTS1-cargos are recognized by the C-terminal tetratricopeptide repeat (TPR) domain of the peroxisomal receptor PEX5 (Gatto *et al*, 2000; Stanley *et al*, 2006). The receptor-cargo complex is tethered to the peroxisomal membrane via its intrinsically disordered N-terminal domain (NTD) (Dammai & Subramani, 2001; Dodt & Gould, 1996; Erdmann & Schliebs, 2005; Rucktaschel *et al*, 2011). At the peroxisomal membrane, PEX5 NTD interacts with the membrane bound components of the translocon PEX14 and PEX13 (**Fig. 1A**) (Elgersma *et al*, 1996; Erdmann & Blobel, 1996; Gould *et al*, 1996; Neufeld *et al*, 2009; Neuhaus *et al*, 2014; Saidowsky *et al*, 2001; Schliebs *et al*, 1999). These interactions may be regulated by weak interactions of PEX5 and PEX14 with peroxisomal membranes (Gaussmann *et al*, 2021; Kerssen *et al*, 2006). After docking, a transient and dynamic pore is formed and the cargo is translocated into the peroxisomal matrix (Erdmann & Schliebs, 2005) (**Fig. 1A**).

**Figure 1.**
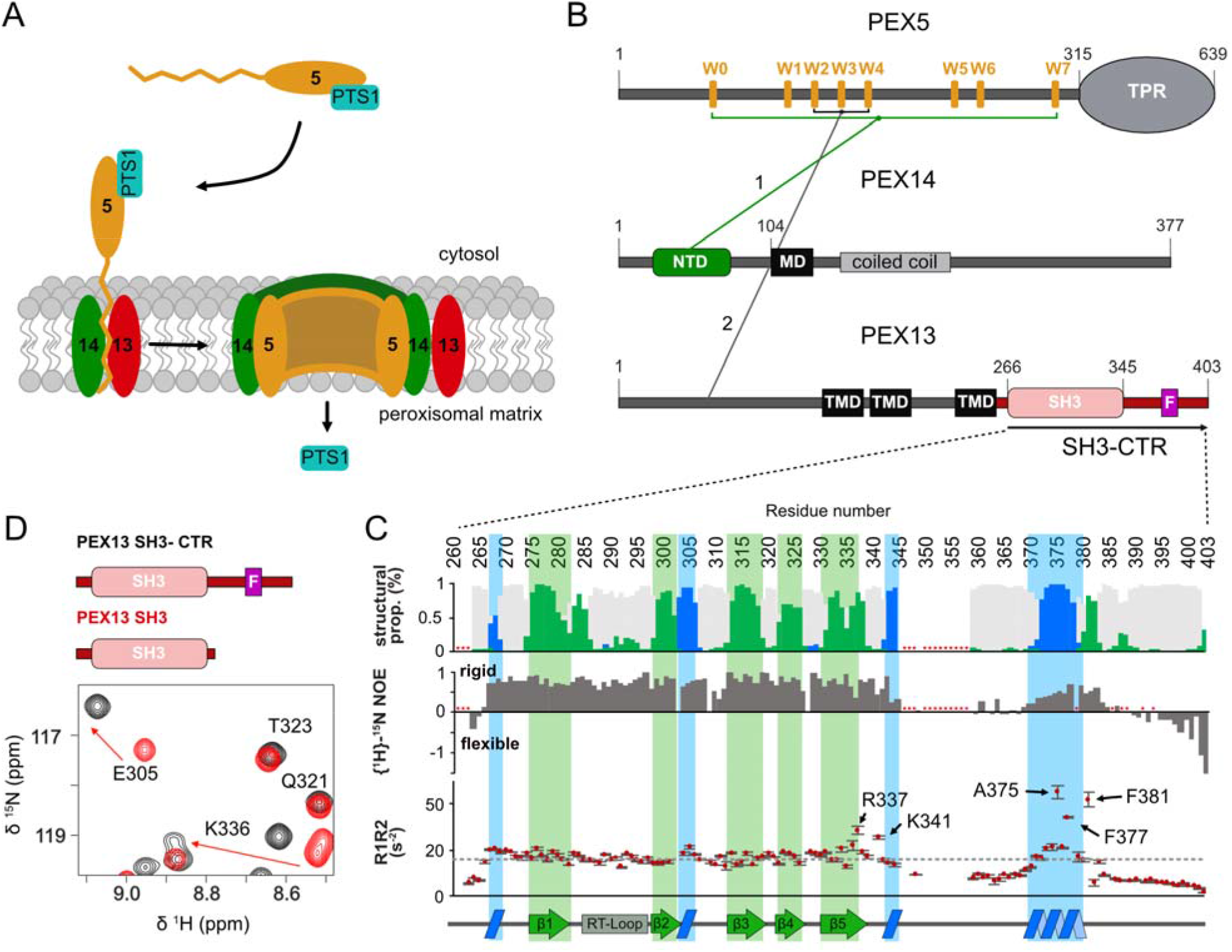
Schematic representation of PTS1 import with interactions the core components and NMR-based analysis of the conformation and dynamics of PEX13 C-terminal region. (A) Schematic overview of cargo recognition, docking and pore formation. (B) Schematic representation of domain architecture and intermolecular interactions of the peroxins PEX5, PEX14 and PEX13 respectively. Lines 1 and 2 between the peroxins indicate known binding events involving the targeted structure elements or motifs (Gaussmann *et al*., 2021; Neufeld *et al*., 2009; Neuhaus *et al*., 2014; Otera *et al*., 2002; Saidowsky *et al*., 2001). (C) Top: ^13^C secondary chemical shifts (Δδ^13^C_α_ - Δδ^13^C_β_) analysed with TALOS+. The propensity for the secondary structure elements random coil, α-helix or β-sheet are represented in grey, blue or green, respectively. Our data support the typical β-sandwich fold of the SH3 domain and the presence of a short α-helical motif comprising the FxxxF motif. Middle: elevated {^1^H}-^15^N heteronuclear NOE values indicate an extended SH3 fold (265-345) and a folded FxxxF motif with similar values to the SH3 domain. Asterisk indicate proline or missing assignment. Bottom: ^15^N *R*_1*_*R*_2_ relaxation rates as a function of amino acid sequence. SH3 core residues (266-335) show an average of 16.6. C-terminal residues R337, K341, A375, F377 and F381 show values of 28.4 ± 1.8, 25.2 ± 0.6, 45 ± 2.6, 34,2 ± 0.6 and 42.0 ± 3.0 respectively. Values higher compared to the average in structured regions indicate the presence of conformational dynamics/and or transient interactions. Secondary structure elements are illustrated at the bottom. (D) Zoom into overlaid spectra of PEX13 SH3-CTR (black) and PEX13 SH3 (red).

Many aspects of peroxisome biogenesis have been studied in yeast, where Pex5/cargo docking is mediated by a membrane-associated complex consistent of the Pex13 SH3, the Pex14 NTD and Pex5 NTD, essential for both PTS1 and PTS2 import (Dodt & Gould, 1996; Elgersma *et al*., 1996; Erdmann & Blobel, 1996; Gould *et al*., 1996; Stein *et al*, 2002). The interactions within the docking complex are mediated (di)aromatic penta peptide motifs (“WxxxF/Y”), of Pex5 with the N-terminal domain of PEX14 and a poly-proline (PxxP) motif of Pex14 that binds to the Pex13 SH3 domain (Barnett *et al*, 2000; Douangamath *et al*, 2002; Erdmann & Blobel, 1995). However, Pex13 might not be directly involved in the translocation process as a purified peroxisome pore *in vitro* comprises only Pex5 and Pex14 (Meinecke *et al*, 2010).

The yeast docking complex differs from the human one by the additional presence of Pex17, which is essential for the peroxisomal import in yeast but seems to be dispensable in humans. Beside this difference, the composition of the human docking complex is like the one in yeast but the homologous interactions are less well studied. Binding between PEX5 (di)aromatic motifs and the globular N-terminal domain of PEX14 (**Fig. 1B, 1**) has been shown to be conserved, but a docking complex with PEX13 similar to the one observed in yeast has not been reported. Human PEX13 is an integral membrane protein with an intrinsically disordered N-terminal domain followed by three transmembrane spans (Barros-Barbosa *et al*, 2019) and a mostly unstructured C-terminal region harboring a SH3 domain. (**Fig. 1D**). PEX13 was first identified and studied in the context of peroxisomal import (Elgersma *et al*., 1996; Erdmann & Blobel, 1996; Gould *et al*., 1996) and Zellweger spectrum disorder (Liu *et al*, 1999). However, our knowledge of PEX13 and its role in peroxisomal import is quite limited. An early study postulated that the PEX5/PEX13 interaction occurs via PEX5 WxxxF/Y motifs W2, W3, and W4 and the PEX13 N-terminal region (**Fig. 1B, 2**), which was shown to be essential for catalase import (Otera *et al*, 2002). Furthermore, a Zellweger mutation W313G located in the PEX13 SH3 domain did not abolish the interaction with PEX14 and was demonstrated to disrupt PTS1 but not PTS2 import (Krause *et al*, 2006; Krause *et al*, 2013).

In this study, we present the first structural characterization of the PEX13 SH3 domain and C-terminal region and identify molecular interactions of PEX13 with PEX5 and PEX14 by combining biophysical methods, structural biology and functional analysis in cells. We show that a novel FxxxF peptide motif in a C-terminal extension of PEX13 mediates an autoinhibitory interaction with the PEX13 SH3 domain. Our binding studies demonstrate that a dynamic network of interactions of the PEX5 NTD with PEX13 and PEX14 modulates peroxisomal protein import.

## Results

### The PEX13 C-terminal region harbors an SH3 domain followed by a FxxxF motif

To study the conformation of the PEX13 C-terminal region, comprising a SH3 domain and a C-terminal unstructured extension (SH3-CTR), we used NMR spectroscopy. The ^1^H-^15^N correlation spectra of the PEX13 SH3-CTR (residues 261-403) shows well dispersed signals and additional signals with narrow linewidth, which correspond to the globular SH3 domain and the disordered C-terminal region, respectively (**Supplementary Fig. 1A**). NMR ^13^C_α_ and ^13^C_β_ secondary chemical shifts are consistent with the secondary structure of SH3 domains composed of five β-strands (β1 to β5) (Saksela & Permi, 2012). An additional α-helical motif located in the C-terminal region (371-383) downstream of the SH3 domain encompasses an FxxxF motif (**Fig. 1C, Top, Supplementary Fig. 1B**), which is highly conserved across mammals (**Supplementary Fig. 1C)**. We then compared NMR correlation spectra of the PEX13 SH3-CTR and SH3 domain. Surprisingly, significant chemical shift differences are seen in the SH3 domain, which map to the regions of β1, β5 and in the region of β2 and an alpha turn. (**Fig. 1D, Supplementary Fig. 1A, D**). Then, we recorded {^1^H}-^15^N heteronuclear NOEs (hetNOE), which reflect the flexibility of the backbone at sub nanosecond timescales. Reduced backbone flexibility indicated by the hetNOE data shows that the PEX13 SH3 fold extends beyond the typical SH3 structural elements compared to the yeast Pex13 and other human SH3 domains (**Supplementary Fig. 1C**). In the C-terminal region, around residue 350, hetNOE values decrease to 0, showing high conformational flexibility of the C-terminal region. However, the FxxxF motif in this region shows hetNOE values of ∼0.8, comparable to those in the globular SH3 domain (**Fig. 1C, middle**), indicating conformational rigidity. This is further supported by the product of NMR ^15^N *R*_1_ and *R*_2_ relaxation rates. While similar values with an average of 16.6 s^-2^, are seen for the rigid core domain, some C-terminal residues of the SH3 domain and the FxxxF motif show significant elevated values ranging from 25 s^-2^ to 45 s^-2^, indicating conformational exchange (**Fig. 1C, bottom)** (Kneller *et al*, 2002). These suggests potential transient intramolecular interactions between the FxxxF motif and the PEX13 SH3 domain.

### The PEX13 SH3 domain is autoinhibited by the C-terminal FxxxF motif

The domain boundaries defined by the NMR analysis were used to generate chimeric constructs which contain the PEX13 SH3 domain and the FxxxF motif (D_371_EQEAA<u>F</u>ESV<u>F</u>V_383_) separated by GGGGS (GS) linkers. Structures of the apo SH3 domain (**Supplementary Fig. 2A**) and SH3-(GS)_2_-FxxxF (**Fig. 2A**) were solved by X-ray crystallography at 1.8Å and 2.3Å resolution, respectively (**Supplementary Table 1**). A comparison of the apo PEX13 SH3 and the complex structure show that the SH3 fold is highly similar (backbone coordinate RMSD = 0.39) (**Supplementary Fig. 2C**). Analysis of the structure of the SH3 domain shows a network of polar contacts between the N- and C-terminal regions stabilizing the β1/β5 interaction (**Supplementary Fig. 2B**), consistent with the extended domain boundaries observed by NMR. The structure of the complex shows interactions of the α-helical FxxxF motif and SH3 domain mediated by hydrophobic contacts of the two phenylalanines, which clamp around β1 and β5 (**Fig. 2A, C**), and polar interactions involving sidechain and backbone contacts. Backbone hydrogen-bonds are formed between A376 and G335 as well as S379, F381 and K336 while negatively charged sidechains E374 and E378 show electrostatic interactions with K304, E305, K336 and a water molecule (**Fig. 2B**). Interestingly, nine out of the eleven Arg and Lys residues are located at the FxxxF binding surface, causing a highly positive charge, which is favorable for binding negative charged peptides such as the C-terminal FxxxF motif (**Fig. 2C**). The PxxP binding site located at the other side of the SH3 domain, is on the other hand mostly negatively charged. The crystal structure was confirmed in solution by NMR titration experiments of the isolated SH3 domain with a FxxxF peptide (350-403) showing strong chemical shift perturbations in the binding site expected from the crystal structure, where the spectrum at saturated binding is very similar to the native SH3-CTR (**Supplementary Fig. 2D, E)**. Furthermore, the static light scattering (SLS) analysis of PEX13 SH3-CTR indicates a molecular weight of 15.6 ± 0.1 kDa, which correlates well with the calculated mass of 15.6 kDa (**Fig. 2D**). This confirms that the FxxxF/SH3 interaction occurs intramolecularly and does not involve oligomerization of the construct at the measured concentration. These results suggest that the PEX13 SH3-CTR adopts an auto-inhibited state, where the C-terminal FxxxF motif interacts with the SH3 domain.

**Figure 2.**
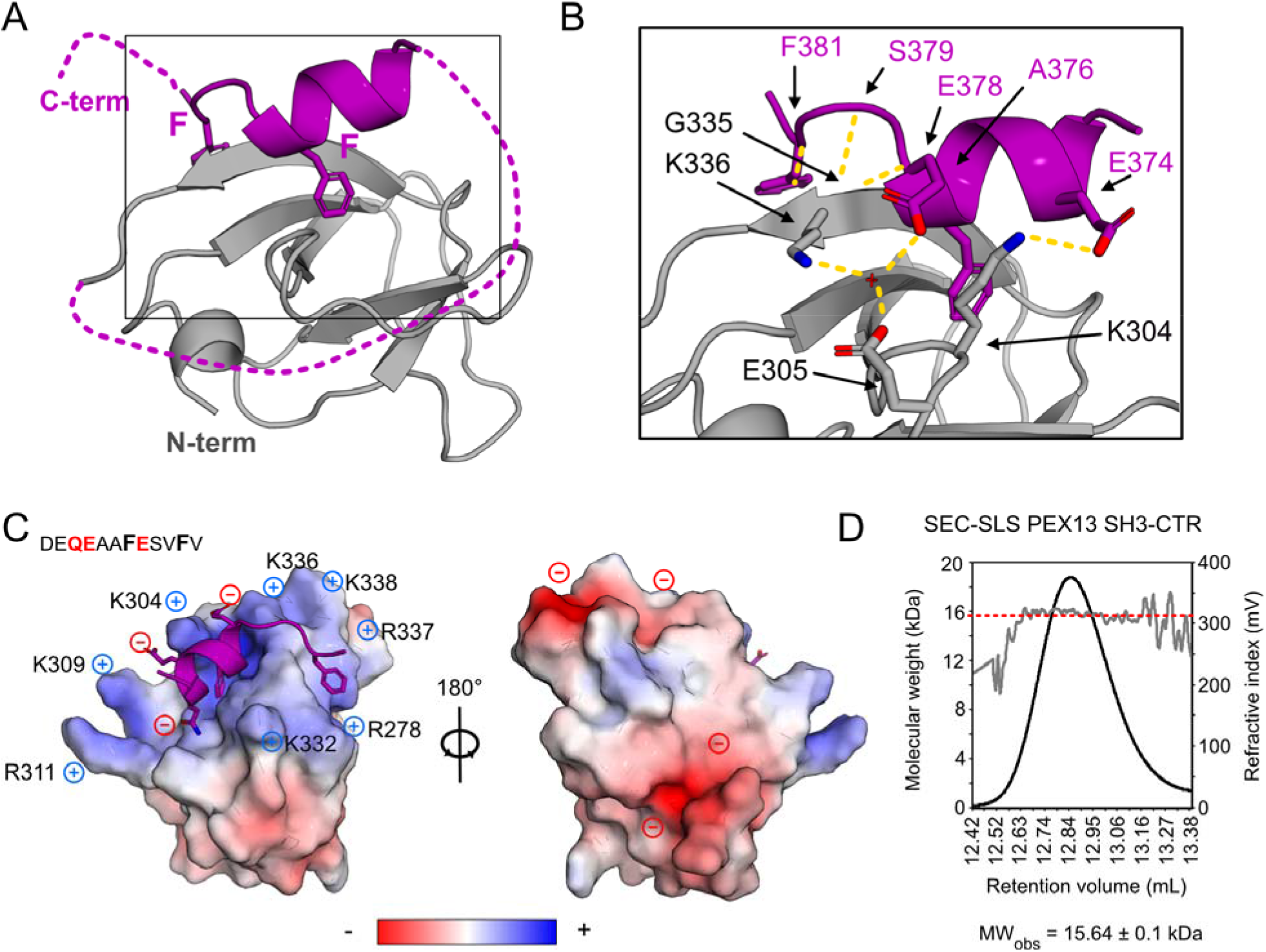
Structural analysis of PEX13 SH3 in complex with FxxxF motif. (A) Crystallographic structure of PEX13 SH3 2GS FxxxF showing the α-helical FxxxF motif, which clamps β1 and β5 between the two Phe residues. (B) Zoom view visualizing the hydrogen bond network between the SH3 domain and the FxxxF peptide. Polar backbone contacts are mediated by A376 and G335 as well as S379, F381 and K336 and sidechains E374 and E378 are coordinated by K304, E305, K336 and a water molecule. (C) Electrostatic surface representation showing the positively charged FxxxF binding site which is caused by seven Arg or Lys residues. The peptide in contrast, is negatively charged which is favored for the binding (Q373, E374 and E387). A 180° rotation on the Y axis reveals a negatively charged backside. (D) Static light scattering analysis of PEX13 SH3-CTR shows the molecular weight (red dashed line) of 15.64 ± 0.1 kDa versus the calculated mass of 15.56 kDa indicating a monomeric state.

### The PEX13 FxxxF motif binds to the PEX14 NTD

We next evaluated interactions of PEX13 with the core components of the import machinery. First, we analyzed the PEX13 SH3-CTR / PEX14 NTD interaction by NMR titrations monitoring effects on the ^15^N labeled PEX14 NTD upon titration of unlabeled PEX13 SH3-CTR. Significant chemical shift perturbations are observed for amide signals (**Fig. 3A, Supplementary Fig. 3A**) and with a profile similar to the known interaction of PEX14 NTD with the PEX19 FxxxF motif (**Fig. 3C**) (Neufeld *et al*., 2009). A sequence comparison of the two motifs including the five preceding residues from PEX13 and PEX19 (**Fig. 3B**) shows strong similarities with four identical and two similar residues (**Fig. 3B**). Not surprisingly, mapping the chemical shift perturbations onto the PEX14 NTD structure highlights the involvement of key residues that are also involved in binding of PEX5 WxxxF/Y motifs (Neufeld *et al*., 2009; Neuhaus *et al*., 2014). The PEX14 NTD / PEX13 FxxxF interaction was analyzed by ITC showing binding affinities for the auto-inhibited PEX13 SH3-CTR (261-403) or a PEX13 FxxxF construct (starting at the linker region just after the SH3 domain, residues 350-403) corresponding to a *K*_*D*_ of 5.4 µM and 2.8 µM respectively (**Fig. 3E, Table 1**).

**Table 1.** Isothermal titration calorimetry of PEX14 NTD with PEX13 SH3-CTR or PEX13 C-terminal peptide (351-403).

| Construct | N | K<sub>D</sub> (μM) | ΔG (kJ/mol) | ΔH (kJ/mol) | -TΔS (kJ/mol) |
| --- | --- | --- | --- | --- | --- |
| PEX13 SH3-CTR | 1 | 5.4 ± 1.2 | -30.2 ± 0.50 | -13.4 ± 1.15 | -16.7 ± 1.47 |
| PEX13 FxxxF | 1 | 2.8 ± 0.4 | -31.8 ± 0.36 | -48.3 ± 1.3 | 16.5 ± 1.62 |

**Figure 3.**
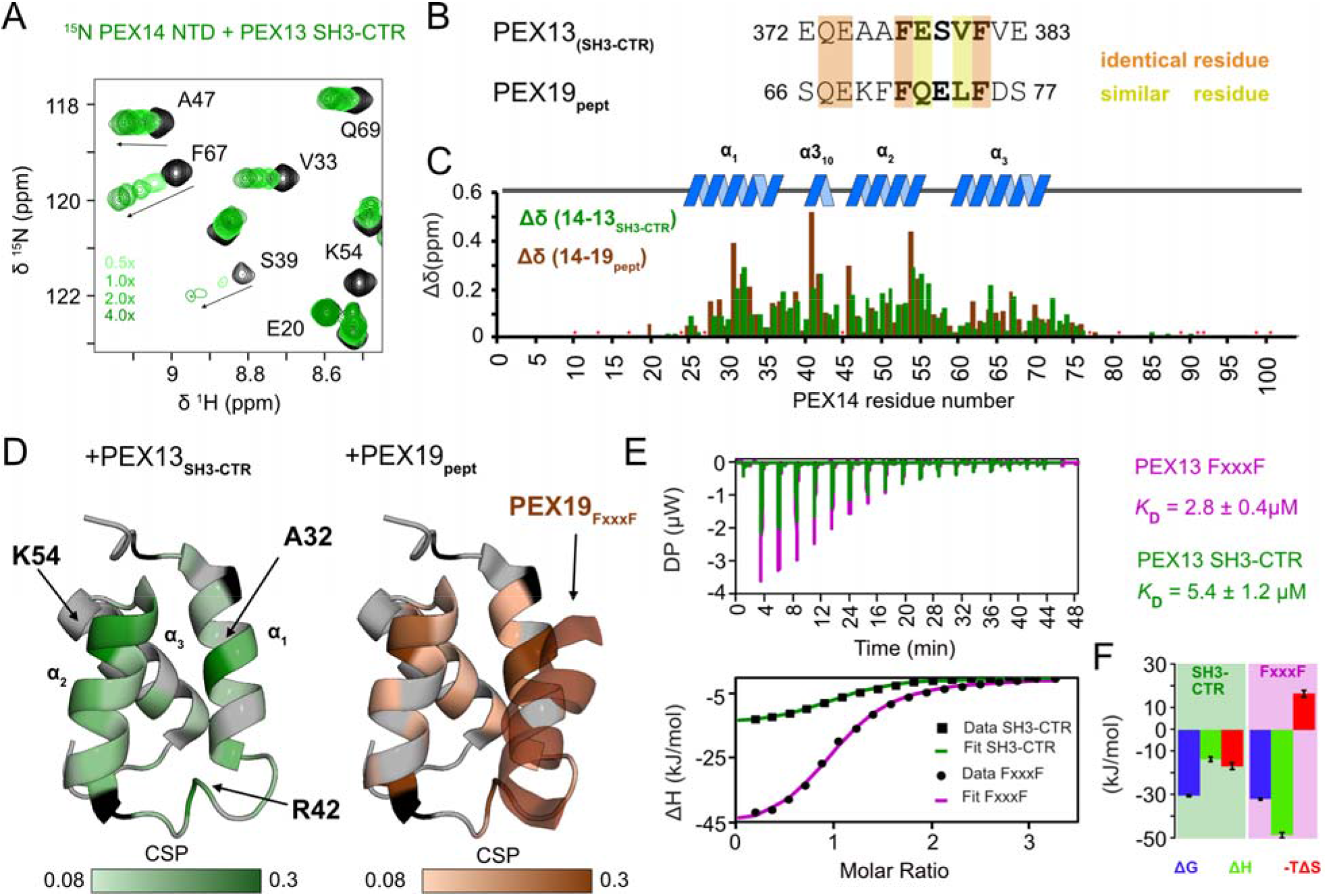
Interaction of PEX14 NTD with PEX13 SH3-CTR in comparison with PEX19 (66-77) (A) ^1^H,^15^N correlation spectra of ^15^N-labelled recombinant PEX14(NTD) free (black), and in complex with PEX13 SH3-CTR (green scale). (B) Sequence alignment of PEX13 and PEX19 FxxxF motifs. Red and yellow boxes indicate identical and similar residues. (C) NMR chemical shift perturbations of PEX14 NTD in the presence of PEX13 SH3-CTR (green) or PEX19 peptide (brown) (Neufeld et. al.) plotted on the sequence with indicated secondary structure elements above. Asterisk indicate proline or missing assignment. (D) Chemical shift perturbations (0.08 to 0.3 ppm) of PEX13 SH3-CTR (left) and PEX19 peptide (right) mapped on the PEX14 NTD/PEX19 66-77 structure (2w85, Neufeld et al.). (E) ITC experiments of PEX14 NTD with PEX13 SH3-CTR (green) and PEX13 FxxxF (pink) showing very different energetics but the same one to one stoichiometry. (F) Energetic contribution of the PEX14 NTD interaction with PEX13 SH3-CTR (left graph) and PEX13 FxxxF peptide (right graph).

Although the dissociation constants (*K*_*D*_) are in a similar range, with two-fold stronger binding for the isolated FxxxF peptide, the underlying energetics are notably different. While the interaction with PEX13 SH3-CTR profits from enthalpic and entropic effects, which likely arises from the transition of the PEX13-associated motif to the PEX14 bound form, the interaction with free FxxxF comes with an entropic penalty, reflecting a free-to-bound transition for the FxxxF motif (**Fig. 3F, Table1)**. The ITC experiments further demonstrate a stoichiometry of 1:1 in both cases (**Fig. 3E lower panel, Table 1)**. A comparison of the interaction strength of the PEX13 FxxxF and the PEX19 FxxxF motif (*K*_*D*_ = 9.2 µM, Neufeld *et al*. (2009)) shows a 3 times stronger binding of PEX13 FxxxF towards PEX14 NTD. Taken together, these data imply that the auto-inhibited state of PEX13 SH3-CTR is readily released upon PEX14 binding. These findings also show that the human SH3 can interact with (di)aromatic peptide motifs on a surface opposite to the PxxP binding region, as has previously been shown for the yeast Pex13 SH3 domain (Douangamath *et al*., 2002).

To assess whether the interaction site in PEX13 is limited to the FxxxF motif or involves additional regions, we titrated unlabeled PEX14 NTD (1-104) onto ^15^N labeled PEX13 SH3-CTR (**Fig. 4**). Interestingly, strong chemical shift perturbations are seen not only for the FxxxF motif but also for residues in the SH3 domain **(Fig. 4A, B, Supplementary Fig. 4A)**. Notably, NMR signals of the SH3 domain shift towards their position in the SH3 domain alone (**Fig. 4B, C yellow boxes**). Signals of the FxxxF peptide (370-386) experience large chemical shift perturbations or line-broadening, down to beyond detection for residues 378 to 383 located in the core motif (**Fig. 4C**). We conclude that the PEX14 NTD interacts solely with the FxxxF motif, and that this interaction releases the auto-inhibited conformation of the PEX13 SH3 domain (**Fig. 4D**).

**Figure 4.**
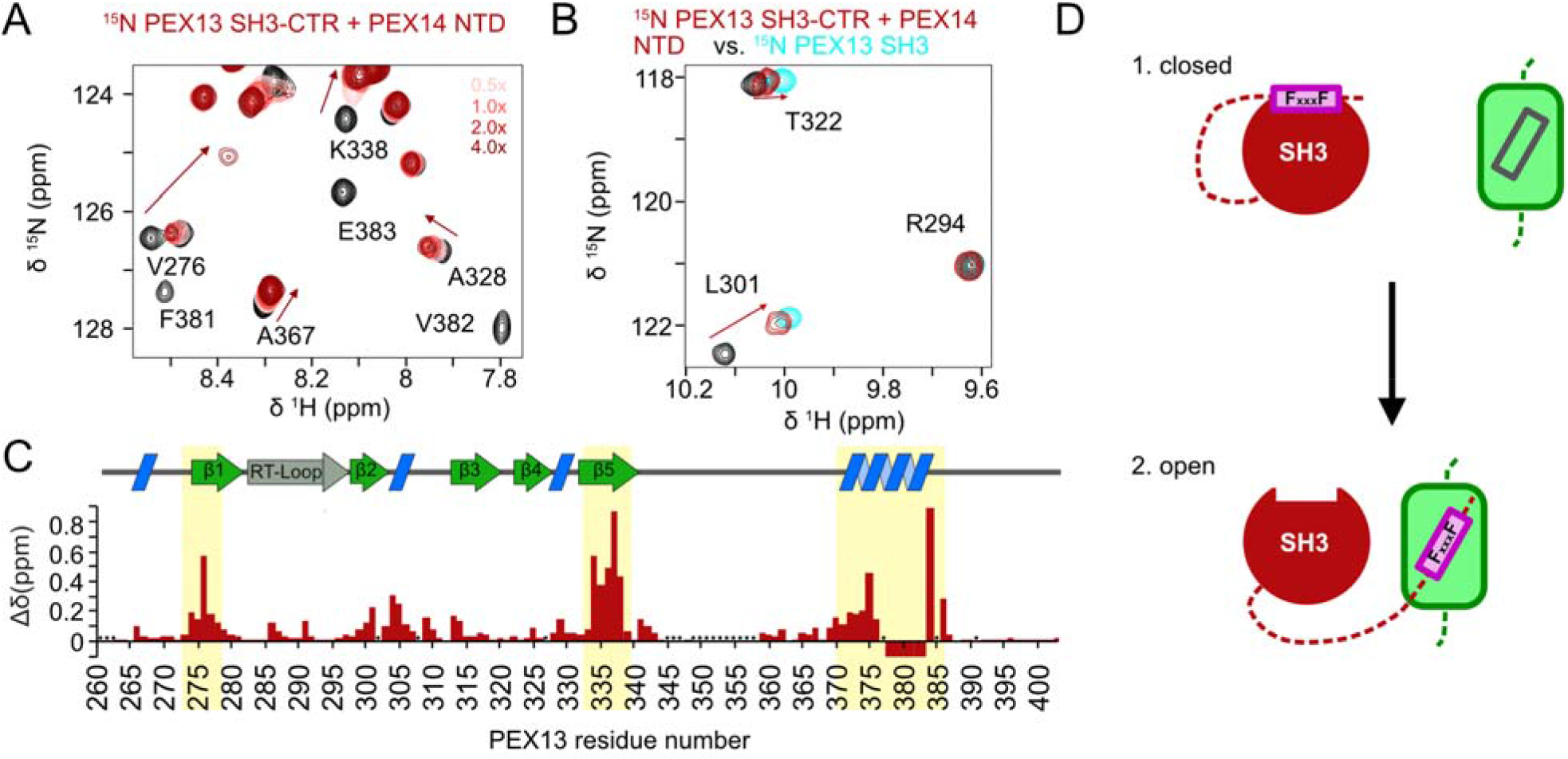
NMR titration of PEX14 NTD onto ^15^N PEX13 SH3-CTR. (A) Spectra overlay from NMR titration of unlabeled PEX14 NTD onto ^15^N labeled PEX13 SH3-CTR where large chemical shift perturbations of resonances from the FxxxF motif were observed. (B) Spectra overlay of free PEX13 SH3-CTR (black), PEX13 SH3-CTR + 4x PEX14 NTD (dark red) and apo PEX13 SH3 (blue) showing the transfer from the closed conformation of the PEX13 SH3-CTR back to apo form of PEX13 SH3. (C) Chemical shift perturbations mapped on the sequence and structural elements (above) of PEX13 SH3-CTR visualizing the effect of the opening on the structural elements β1, β5 and the FxxxF motif. Signals that experience large chemical shift perturbations or line-broadening down to beyond detection are indicated with negative values (D) Schematic representation of the opening process.

### PEX13 SH3 interactions with poly-proline motifs

In yeast, the Pex14/Pex13 interaction is mediated by a class II poly-proline motif of Pex14, which binds to the Pex13 SH3 domain (Douangamath *et al*., 2002). Human PEX14 also harbors a PxxP motif downstream of its NTD (residues 87-102), similar to that in yeast (residues 85-94). NMR titrations with unlabeled PEX14 NTD_long_ (1-113) onto ^15^N labeled PEX13 SH3 were used to evaluate this potential interaction. No significant chemical shift perturbations were observed, showing that this interaction is not conserved from yeast to human (**Supplementary Fig. 4B**). Our results are in agreement with previous experiments using co-immunoprecipitation that showed an interaction of PEX14 with the PEX13 Zellweger mutant W313. This mutation is located in the SH3 domain and destabilizes the SH3 fold which affects multiple interactions of the SH3 domain. The mutation did not affect the human PEX14/PEX13 interaction, while the same mutation abolished Pex14 binding in the yeast homologues (Krause *et al*., 2013). We thus investigated whether human PEX13 SH3 has any ability to bind PxxP motifs. Interestingly, PxxP motifs present in the N-terminal region of PEX13 showed significant binding in NMR titration experiments with the PEX13 SH3 domain and the PEX13 SH3-CTR. These experiments demonstrate that, in principle, the human SH3 domain can mediate PxxP interactions and that these are independent, and non-overlapping, with the FxxxF binding (**Supplementary Fig. 4C, D)**. However, chemical shift perturbations mapped onto a structural model of a class II PEX13 SH3/PxxP complex indicate an imperfect fit in the second proline binding pocket occupied with an isoleucine, suggesting that the human PEX13 SH3 domain shows interactions with class I PxxP peptides, distinct from the yeast orthologue (**Supplementary Fig. 4F)**.

### PEX5 WxxxF/Y motifs compete with the internal FxxxF motif on PEX13 SH3

We then characterized the molecular interactions between PEX13 and PEX5. The PEX5 NTD (1-315) but not PEX5 TPR (315-639) domain was found to bind to PEX13 SH3-CTR and PEX13 SH3 (**Supplementary Fig. 5**). Using NMR titrations we show that the eight (di)aromatic peptide motifs of PEX5, also known as W-motifs, bind to the PEX13 SH3 domain or the PEX13 SH3-CTR. Noteworthy, the PEX5 W binding site overlaps with the FxxxF binding and thus competes with the internal motif (**Fig. 5A,B,C, Supplementary Fig. 6A, B**). The PEX5 W4 motif was identified as the strongest binder followed by W2 and W3 (**Fig. 6B**). These PEX5 motifs can compete with the PEX13 internal FxxxF motif, while other (di)aromatic peptide motifs only bind to PEX13 SH3 with low affinity and not to PEX13 SH3-CTR (**Fig. 6C)**. PEX5 W3 induces small CSPs but significant line-broadening on PEX13 SH3 (**Fig. 6B star**), which indicates strong binding as well. These observations are supported by ITC experiments, which show *K*_*D*_’s of 43, 88, and 102 µM for W4, W2 and W3, respectively. To relate these affinities, we evaluated the binding of the PEX13 internal FxxxF motif in *trans*, which shows a *K*_*D*_ of 27 µM **(Table 2)**, and thus stronger than any of the (di)aromatic peptide motifs of PEX5. Affinities of the other motifs were too weak to be measured accurately by ITC (**Fig. 5. D, Supplementary Fig. 6 C, D, Table 2)**. Binding of the same PEX5 motifs to PEX14 NTD is overall much stronger, ranging from 21 nM to 3136 nM (**Fig. 5D, E, Supplementary Fig. 7, Table 3**). These values are in agreement with previous studies in different buffer conditions (Gopalswamy *et al*, 2022). Of note, amongst the eight (di)aromatic peptide motifs in PEX5, W4 shows the highest relative binding affinity for PEX13 SH3 and the weakest interaction with the PEX14 NTD (**Fig. 4E, 5D, E**). To investigate the binding mode of PEX5 W4 to PEX13 SH3 in more detail, we crystallized a PEX13 SH3 GS W4 chimera and solved the structure by X-ray crystallography at 2.3 Å (**Fig. 5F, Supplementary Table 1**). In contrast to the internal FxxxF motif, the binding interface is limited to the core motif driven by hydrophobic interactions and few hydrogen bonds from R183, Y185 and Y188 to the backbone or K304 sidechain **(Fig. 5G**). Nevertheless, polar backbone interactions involving G335 and K336 as well as coordination from K304 seem to be important since they are conserved from PEX13 FxxxF to PEX5 W4 (**Fig. 5H, colored lines**). These results show that all (di)aromatic penta peptide motifs from PEX5 NTD are able to bind the isolated PEX13 SH3 domain but only W4, W2 and W3 can compete with the internal FxxxF motif. The structure of the PEX13 SH3 - W4 complex shows a limited binding interface that lacks the electrostatic interactions that are seen with the PEX13 FxxxF motif (**Fig 2**).

**Table 2.** Isothermal titration calorimetry of PEX13 SH3 with PEX5 W2, W3, W4 or PEX13 C-terminal peptide (351-403) with WT FxxxF or introduced PEX4 W4 motif.

| Peptide | N | K<sub>D</sub> (μM) | ΔG (kJ/mol) | ΔH (kJ/mol) | -TΔS (kJ/mol) |
| --- | --- | --- | --- | --- | --- |
| PEX5 W2 | 1 | 88.1 ± 0.1 | -23.17 ± 0.04 | -27.00 ± 0.47 | 3.81 ± 0.47 |
| PEX5 W3 | 1 | 102.2 ± 9.9 | -22.80 ± 0.20 | -14.65 ± 3.25 | -8.14 ± 3.46 |
| PEX5 W4 | 1 | 43.4 ± 4.4 | -24.93 ± 0.24 | -37.30 ± 2.66 | -12.38 ± 2.95 |
| PEX13 FxxxF | 1 | 26.9 ± 3.7 | -26.13 ± 0.38 | -24.73 ± 1.38 | -1.42 ± 1.69 |
| PEX13 FxxxF to W4 | 1 | 6.4 ± 0.2 | -29.67 ± 0.09 | -53.50 ± 1.53 | 23.80 ± 1.47 |

**Table 3.** Isothermal titration calorimetry of PEX14 NTD with PEX5 W-peptide motifs.

| Peptide | N | K<sub>D</sub> (nM) | ΔG (kJ/mol) | ΔH (kJ/mol) | -TΔS (kJ/mol) |
| --- | --- | --- | --- | --- | --- |
| PEX5 W0 | 1 | 55.1 ± 0.1 | -41.53 ± 0.44 | -46.43 ± 2.56 | 4.91 ± 3.00 |
| PEX5 W1 | 1 | 72.3 ± 6.4 | -40.83 ± 0.24 | -77.27 ± 2.89 | 36.43 ± 2.64 |
| PEX5 W2 | 1 | 73.6 ± 5.9 | -40.96 ± 0.18 | -49.10 ± 1.13 | 8.33 ± 1.23 |
| PEX5 W3 | 1 | 136.5 ± 34.3 | -39.33 ± 0.62 | -76.00 ± 1.27 | 36.63 ± 0.69 |
| PEX5 W4 | 1 | 3136.7 ± 417.8 | -31.50 ± 0.33 | -55.17 ± 2.36 | 23.70 ± 2.00 |
| PEX5 W5 | 1 | 20.5 ± 3.3 | -43.96 ± 0.38 | -74.47 ± 1.09 | 30.47 ± 1.49 |
| PEX5 W6 | 1 | 636.3 ± 20.22 | -35.40 ± 0.07 | -73.13 ± 0.58 | 37.70 ± 0.60 |
| PEX5 W7 | 1 | 236.0 ± 21.33 | -37.90 ± 0.20 | -64.63 ± 1.22 | 26.77 ± 1.31 |

**Figure 5.**
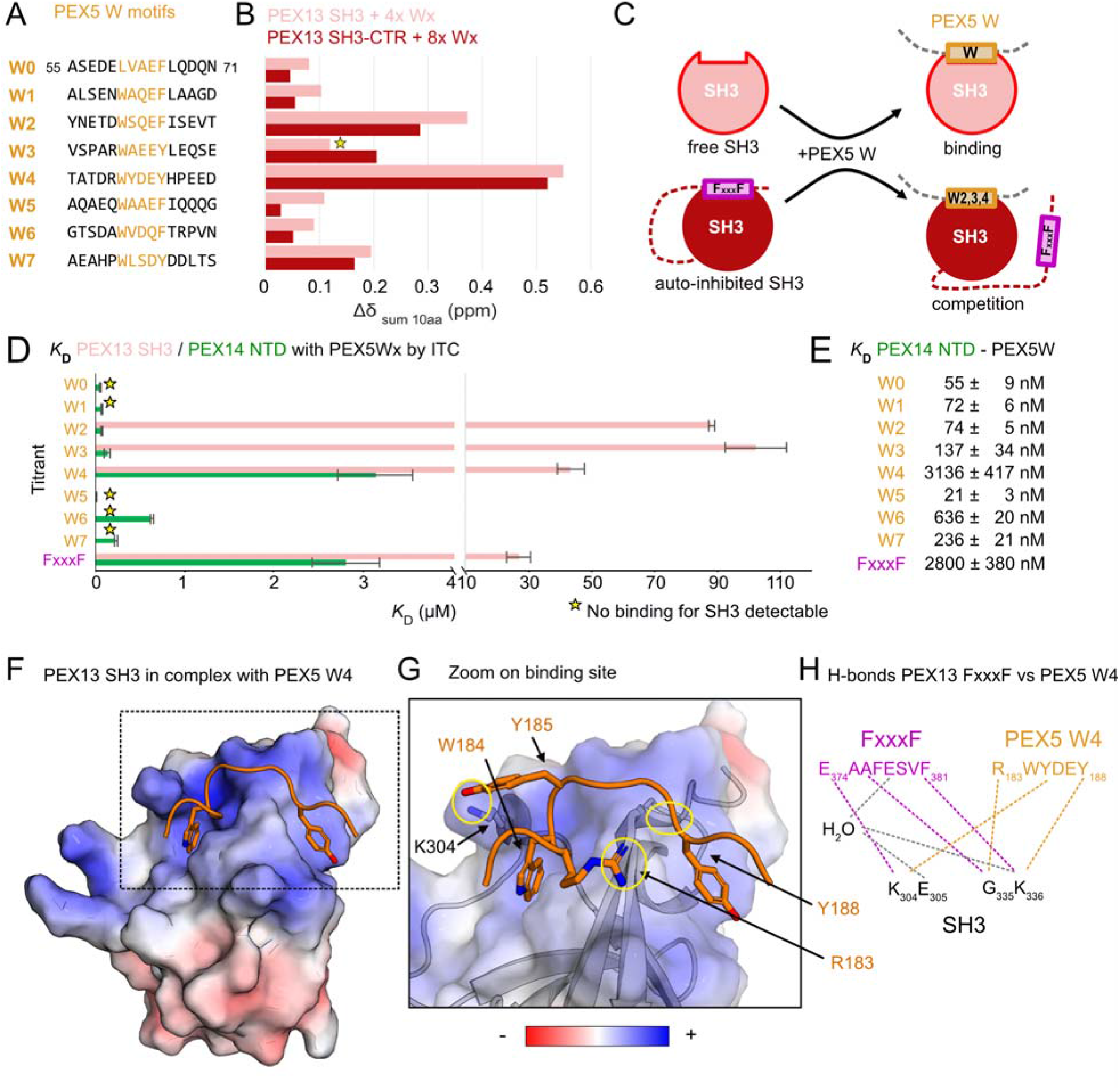
Interaction of PEX5 (di)aromatic peptide motifs with PEX13 SH3 or SH3-CTR. (A) Overview of PEX5 (di)aromatic peptide motifs. W0 was expressed as PEX5 1-76 while other W motifs were purchased as peptides as listed. (B) Induced Chemical shifts changes of PEX13 SH3 or SH3-CTR upon addition of 4x or 8x PEX5 (di)aromatic peptide motifs represented as the sum of 10 involved residues. The star indicates W3 which shows less chemical shift perturbation but extensive line-broadening. (C) Schematic representation of PEX13 SH3 / W peptide binding (top) or PEX13 SH3-CTR / W peptide competition. (D) Plot of triplicate *K*_D_ values from ITC experiments of PEX13 SH3 or PEX14 NTD with PEX5 W and FxxxF peptide motifs. (E) *K*_D_ values from ITC experiments in numbers. (F) Electrostatic surface representation of the PEX13 SH3 GS W4 structure at 2.3 Å. (G) Zoomed view of the W binding site showing polar interactions marked with yellow circles (H) Schematic representation of the conserved hydrogen-bond network of PEX13 SH3 / FxxxF (pink) and PEX5 W4 (orange) interaction. Additional contact sites are marked in gray.

**Figure 6.**
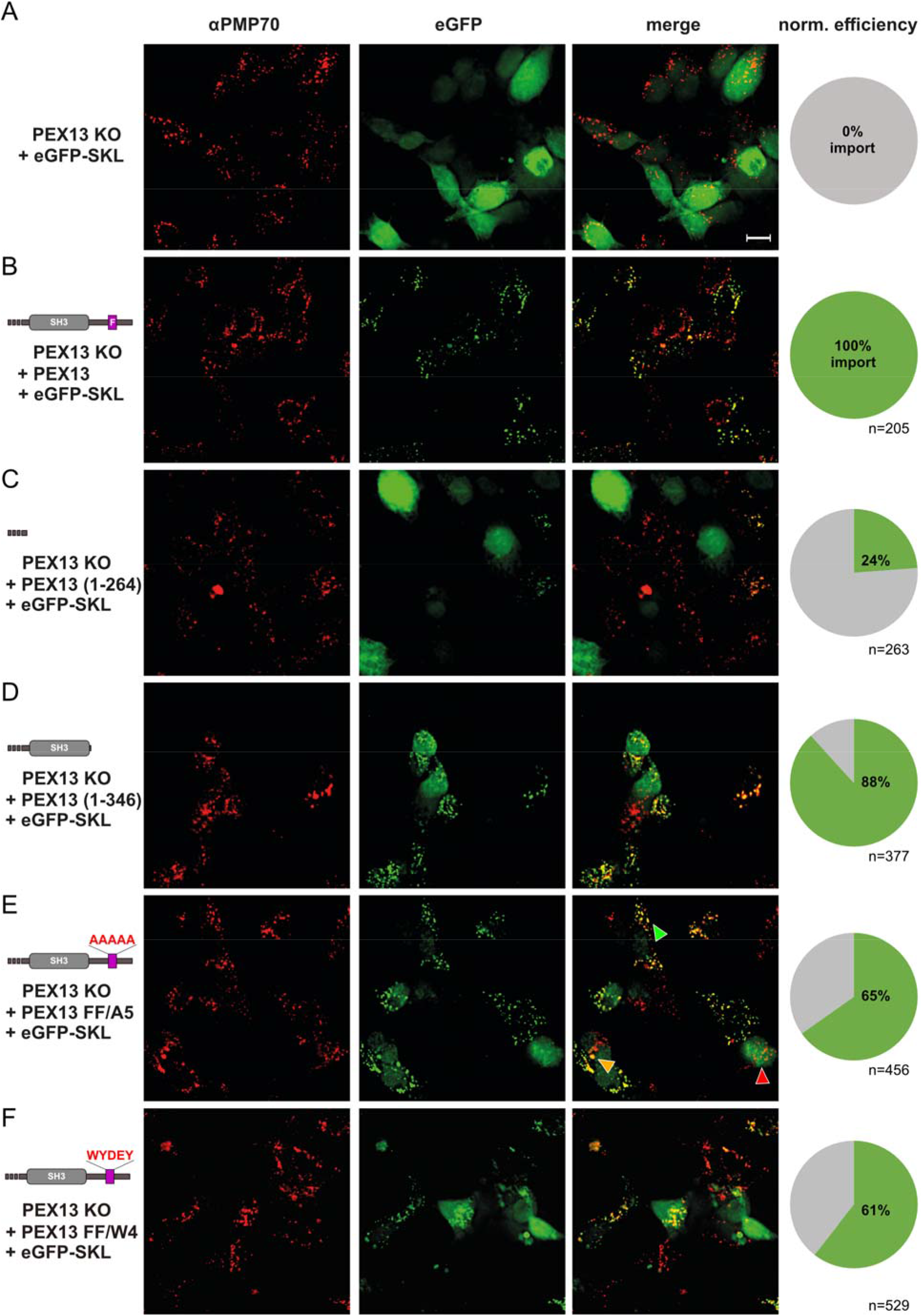
PEX13 SH3-proximal FxxxF motif modulates PTS1 import. PEX13-deficient T-REx cells were transfected with different PEX13 truncation and mutation variants as indicated on the left side in (A-E) to monitor the rescue of PTS1 import defect via fluorescence microscopy. Detection of peroxisomal membranes was achieved via PMP70 specific antibody indicated as red congruent punctate patterns (left panels). Scale bar: 10 µm (A) Expression of solely eGFP-SKL did not rescue PTS1 import indicated by a diffuse cytosolic green eGFP signal (middle and right panels) and served as negative control. (B) Transfection of the cells with PEX13 FL located on a bicistronic vector together with eGFP-SKL led to a congruent punctate pattern colocalizing with PMP70 indicating rescued import. Import efficiencies of other PEX13 variants were normalized to PEX13 FL. (C) Truncation of the full C-terminus (1-264) had a large effect on the function of PEX13 in peroxisomal import, which was seen only for 24% of the cells. (D) Expression of the truncated PEX13 1-346 showed an import efficiency of 88%, whereas expression of PEX13 FF to A5 (E) and FF to W4 (F) reduced the rescue efficiency to 65% and 61%, respectively. Besides the phenotype of import (E, green arrow) and non-import (E, red arrow), some cells showed a partial import (E, orange arrow) indicated by diffuse and punctuate eGFP signal.

### The PEX13 FxxxF motif modulates PTS1 import

The functional significance of the PEX13 SH3-CTR with the modulation of the SH3 domain by the proximal FxxxF motif was addressed using a cellular import complementation assay. Here the complementing activity of PEX13 variants was analyzed in a cell-based model using T-REx PEX13 KO cells with non-functional PTS1 import generated by CRISPR/Cas9 genome engineering. The PEX13-deficient cells were transfected with a bicistronic vector encoding for i) eGFP-SKL as marker for peroxisomal PTS1 import and ii) full-length PEX13 or PEX13 truncation and mutation variants.

Cells solely expressing eGFP-SKL showed no import activity indicated by the diffuse distribution of eGFP in the cytosol (**Fig. 6A**). In contrast, transfection of PEX13 FL results in functional import represented by the congruent punctate pattern of eGFP and co-localization with the peroxisomal marker protein PMP70 (**Fig. 6B)**. Besides functional and non-functional import, cells with partial import with diffuse and punctate eGFP pattern were observed. These cells were counted as import-competent, as they were capable of importing at least some of the eGFP-SKL (**Fig. 6E**). We first assessed the importance of the full C-terminal region (SH3-CTR). The re-introduction of a PEX13 truncation (1-264) lacking the SH3 domain and C-terminal region reduced the import capability drastically to 24% **(Fig. 6C)** compared to wild-type **(Fig. 6B)**. We further investigated the role of the intrinsic FxxxF motif by truncation or mutation. Interestingly, transfection of PEX13 KO cells with the PEX13 1-346 truncation lacking the FxxxF motif, showed a restored import of 88% (**Fig. 6D)**. Substitution of the FxxxF motif by polyA (FF/A5), which was supposed to have non or reduced inhibition ability, led to an import efficiency of 65% (**Fig. 6E)**. Noteworthy, no interaction between the SH3 domain and a C-terminal peptide harboring this FxxxF to A5 mutation was observed by NMR titrations (**Supplementary Fig.8A**). Complementation experiments with the PEX13 FF to W4 mutant, which was shown to bind stronger to the PEX13 SH3 domain in trans as the wild-type with 6µM (W4 core mutation) compared to 27µM (FxxxF) (**Supplementary Fig.8B, Fig. 5D)**, showed a total import efficiency of 61 % **(Fig. 6F)**. Note that in this experiment only the core motif (<u>F</u>ESV<u>F</u>) was replaced with W4 (<u>W</u>YDE<u>Y</u>) which is different to the ITC experiments in figure 5, where the full 15mer sequence of PEX5 W4 was used. The exchange of only the core motif retains the electrostatic interactions of the upstream negatively charged sequence with the positively PEX13 SH3 binding interface and combines it with the stronger hydrophobic interactions of W4 motif indicated by a larger enthalpic contributions (**Table 2)**.

These results highlight the importance of the PEX13 C-terminal region including the SH3 domain and the FxxxF motif as regulatory element in PTS1 import. Our cellular assays demonstrate a drastic effect on import upon deletion of the C-terminus (SH3-CTR) and a modulation of import by the SH3-proximal FxxxF motif.

### The PEX13 FxxxF motif modulates PEX5 interactions in cells

We further verified the presence of the newly identified interactions between PEX13, PEX5 and PEX14 in immunoprecipitation experiments using the previous mentioned T-REx PEX13 KO cells with re-introduced PEX13 variants. We tested PEX13-FL, the C-terminal truncation, which ends after the SH3 domain (PEX13 1-346) (**Fig. 7A**) and the two FxxxF mutants; FxxxF to AAAAA (FF/A5) and FxxxF to W4 (**Fig. 7B**). The Immunoprecipitation experiment was set up to pull down cellular PEX5 by a PEX5 antibody. Experiments with re-introduced PEX13 full-length (FL), revealed visible amounts of PEX13 (**Fig. 7C, second lane IP**), which were further increased in lysates with the PEX13 1-346 truncation (**Fig. 7C, third lane IP, D**) and the PEX13 FxxxF to A5 mutation (**Fig. 7C, fourth lane IP, D**). Lower amounts of PEX13 were detected for PEX13 harboring the FxxxF to W4 mutation compared to FL (**Fig. 7C, fifth lane IP, D**). Noteworthy is the detection of PEX14 in all samples (**Fig. 7C**). The controls with PEX5 KO /PEX13 FL and PEX5 FL/PEX13 KO show no endogenous level of PEX5 or PEX13, respectively. The load was controlled by detection of GAPDH (**Fig. 7C, lower panel**). Repetitions of this experiment can be seen in **Supplementary Fig. 9**. Overall, the data show that the absence of a functional FxxxF motif increase PEX5 binding while the presence of a stronger WxxxF motif at the C-terminus hampers the PEX5-PEX13 interaction

**Figure 7.**
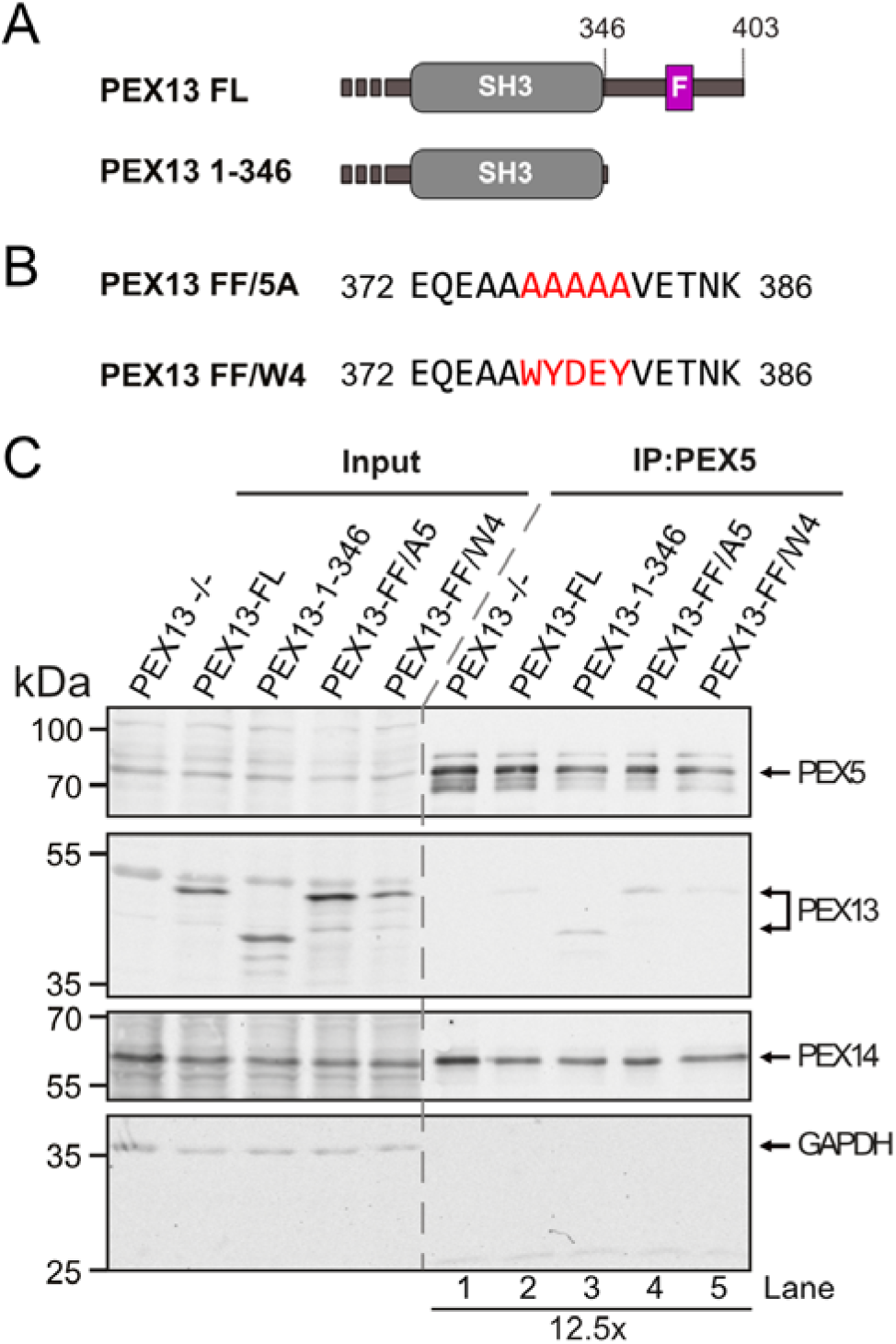
PEX13 FxxxF motif modulates PEX5 binding. PEX13 FL, PEX13 1-346 (A) and the two mutations FxxxF to A5 and FxxxF to W4 (B) were expressed in T-REx PEX13 KO cells. The cell lysates were subjected to immunoprecipitation with PEX5 antibody and analyzed by immunoblotting (C).

## Discussion

Here, we present a comprehensive structural and biophysical analysis of the C-terminal region of human PEX13 and its role in peroxisome biogenesis and function. We show that the PEX13 SH3 domain is essential for PTS1 import and discover the presence of an FxxxF peptide motif in the C-terminal region that mediates an autoinhibitory interaction with the SH3 domain. We observe a novel interaction of the human PEX13 SH3-CTR with the PEX14 NTD that is mediated by the PEX13 FxxxF motif. Importantly, we demonstrate that the SH3 domain of human PEX13 shows binding to (di)aromatic peptide motifs in the PEX5 NTD, similar to yeast, demonstrating evolutionary conservation of these interactions. Moreover, these interactions are further modulated by the autoinhibitory interaction of the proximal FxxxF motif with the PEX13 SH3 domain. Deletion of the internal FxxxF motif decrease of PTS1 import efficiency, indicating that the modulation of the PEX5, PEX13 and PEX14 interactions by the FxxxF motif plays an important role in fine-tuning peroxisome biogenesis and/or matrix protein import in peroxisomes.

SH3 domains commonly recognize PxxP peptide motifs to mediate protein-protein interactions of adaptor proteins (Birge *et al*, 1996) in signal transduction (Schlessinger, 1994). The yeast Pex13 SH3 domain indeed recognizes a class II PxxP motif in Pex14 with the canonical PxxP binding site (Barnett *et al*., 2000; Bottger *et al*, 2000; Douangamath *et al*., 2002; Emmanouilidis *et al*, 2016). Strikingly, this interaction is not conserved from yeast to human as we did not observe an interaction of PxxP motifs in human PEX14 with the PEX13 SH3 domain **(Supplementary Fig. 4B)**. Our results imply that the interaction of the human PEX13 SH3 domain with poly-proline peptide motifs may be distinct, and involves binding to class I peptides, rationalizing why no binding with the class II PxxP motif in the N-terminal region of PEX14 is observed (**Supplementary Fig. 4C,F)**.

The binding of human PEX13 with PEX14 does not involve an SH3 domain/PxxP interaction but is instead facilitated by binding of the PEX13 FxxxF motif to the PEX14 NTD. Not unexpected, the recognition of the PEX13 FxxxF motif by the PEX14 NTD is structurally similar to the interaction with the (di)aromatic PEX5 peptide ligands and an FxxxF motif in the peroxisomal membrane protein transport factor PEX19 (Neufeld *et al*., 2009). This finding is in line with previous studies from Itoh & Fujiki (2006), who mapped the interactions of PEX13, PEX5 and PEX19, to the PEX14 NTD (residues 21-70).

Interestingly, the yeast Pex13 SH3 domain has been shown to also recognize (di)aromatic peptide motifs, using a binding surface opposite to the canonical PxxP binding site (Barnett *et al*., 2000; Bottger *et al*., 2000; Douangamath *et al*., 2002). An interaction of (di)aromatic peptide motifs to other SH3 domains has not yet been reported so far. As residues in the binding site of yeast Pex13 SH3 to Pex5 WxxxF/Y are poorly conserved in human PEX13, it was unclear whether a similar interaction is also possible for human PEX13 **(Supplementary Fig. 1D**). Our results demonstrate that the PEX13 SH3 domain indeed binds to (di)aromatic peptide motifs in PEX5 and that this interaction is conserved from yeast to human. Our crystal structure of PEX13 SH3 with the WxxxF motif reports for the first time a high-resolution details for this non-canonical binding interface of an SH3 domain fold.

A notable difference between the yeast and human PEX13 is that the interaction with (di)aromatic peptide motifs is modulated by the autoinhibition mechanism of PEX13 with the human-specific PEX13 FxxxF motif. Noteworthy, a previous study mapped the PEX13 interaction with WxxxF/Y motifs from PEX5 and to the N-terminal region in PEX13 rather than the SH3 domain (Otera *et al*., 2002). Later, Krause *et al*. (2013) proposed that the PEX13 N-terminal region mediates homo-oligomerization. These observations might be reconciled with recent reports suggesting that the N-terminal region of yeast Pex13 can undergo liquid-liquid phase separation (Gao *et al*, 2022; Ravindran *et al*, 2022), which may thereby indirectly affect the molecular interactions of the C-terminal SH3-CTR module with (di)aromatic peptide motifs.

The network of interactions identified in our study shows that PEX13, PEX5 and PEX14 interactions are modulated by binding of (di)aromatic peptide motifs with overlapping and partially competing binding sites. Thus, the formation of a ternary complex of these three peroxins as reported for yeast is not possible. This suggests an evolutionary increased complexity, where functional activity is not regulated by distinct binding motifs but by multivalency, relative affinity and avidity effects, consistent with the increasing abundance of (di)aromatic peptide motifs from three to eight in yeast and human PEX5, respectively (Kerssen *et al*., 2006; Otera *et al*., 2002; Saidowsky *et al*., 2001). The distinct binding affinities between PEX13/PEX14 **(Fig. 4)**, PEX13/PEX5 and PEX14/PEX5 (**Fig. 5**) suggest a potential sequential binding model. The strong affinity of the PEX5 NTD towards PEX14 NTD may be required for initial docking of cargo-loaded PEX5 at the peroxisomal membrane. This may involve removal of co-factors that might be bound to the PEX5 NTD and thereby restrict accessibility of PEX5 W-motifs other than W0, has been shown to be essential for targeting of PEX5 to the peroxisome (Neuhaus *et al*., 2014; Reglinski *et al*, 2022; Skowyra & Rapoport, 2022). The PEX14/PEX5 interaction may be later replaced by an interaction of PEX13 FxxxF/PEX14 NTD and PEX13 SH3/PEX5 NTD. This would require a high molecular excess of PEX13 considering the lower binding affinity towards PEX5 and PEX14, and is in agreement with the observation of a high molecular mass complex containing only PEX13 or PEX13/PEX14 from previous studies (Gao *et al*., 2022; Itoh & Fujiki, 2006; Ravindran *et al*., 2022; Reguenga *et al*, 2001). It was proposed that PEX14 may change interacting partners depending on the molecular size complexes of PEX14 which are disassembled in the presence of cargo-free PEX5 (Itoh & Fujiki, 2006). Handing PEX5 from the PEX14 NTD over to the PEX13 SH3 could occur during two distinct steps of peroxisomal import; i) as part of the docking event before cargo translocation into the peroxisomal matrix, or ii) to enhance cargo release inside the lumen. Involvement in docking would require that the SH3 domain facing the cytosol, as it was first proposed by Gould *et al*. (1996). However, more recent studies from rat liver indicate that the PEX13 SH3 domain is located inside the peroxisome (Barros-Barbosa *et al*., 2019; Reguenga *et al*., 2001), which would be consistent with the cargo release hypothesis. This hypothesis also implies that the PEX14 NTD faces the cytosol for the docking of PEX5, which is controversial discussed in the field. Recent studies employing proteinase K assays with peroxisomes isolated from rat liver or Xenopus eggs locate the N-terminal domain of PEX14 in the peroxisomal matrix (Barros-Barbosa *et al*., 2019; Skowyra & Rapoport, 2022). Both studies share the same experimental setup, where the cytosolic environment is removed in the step of isolation and thus import or export activity depleted. It is possible that PEX14 NTD is co-translocated with the PEX5-cargo complex into the lumen and after cargo-release and removal of PEX5 flipped back to the cytosol. Considering observations from Otera *et al*. (2002) and Ravindran *et al*. (2022), PEX13 could even act in docking via its N-terminal domain and in cargo release via its C-terminal domain.

Altogether, the results presented in this study together with current knowledge in the field allows us to propose a simplified mechanistic model for the role of PEX13 in peroxisomal matrix import (**Figure 8**). First, a PEX5-cargo-cofactor complex docks by binding to the PEX14 NTD facing the cytosol, with the high affinity W0 motif (**Figure 8A)**. Subsequently, a transient pore consisting of PEX5, PEX14 and PEX13 may be formed, which could involve liquid-liquid phase separation of these components at the membrane (Gao *et al*., 2022; Ravindran *et al*., 2022). The PEX5 TPR domain with the bound cargo is then translocated into the peroxisomal matrix. In the same step, the PEX14 NTD, which is still bound to PEX5 might be translocated to the matrix as well (**Figure 8B**). Next, the PEX5/PEX14 interaction may be replaced by the FxxxF motif of PEX13 assuming a large excess of PEX13, which indeed forms higher homooligomers at the membrane. Concomitantly, the FxxxF motif is released from the PEX13 SH3 domain allowing the transient binding of PEX5. PEX5, PEX14 and PEX13 now form a transient complex where PEX14 is saturated with PEX13 and PEX5 is loosely bound to PEX13 (**Figure 8C, D**). From this intermediate, PEX5 is handed over to the export/ring finger complex consisting of PEX2, PEX10 and PEX12 (Feng *et al*, 2022). With the export of PEX5, the cargo is released into the peroxisomal matrix and PEX14 NTD flips back to the cytosolic side of the membrane via an unknown mechanism (**Figure 8E**). Our model explains the sequential use of overlapping binding sites between PEX5, PEX14 and PEX13 and resolves the apparent inconsistent observations concerning the topology of PEX14. The model further explains why we detect a strong effect on the PTS1 import efficiency when the full SH3-CTR and smaller effects when the FxxxF motif is mutated. With deletion of the SH3, the cycle will be stalled in state B. Mutation or deletion of the FxxxF motif can be compensated by a larger excess of PEX13.

**Figure 8.**
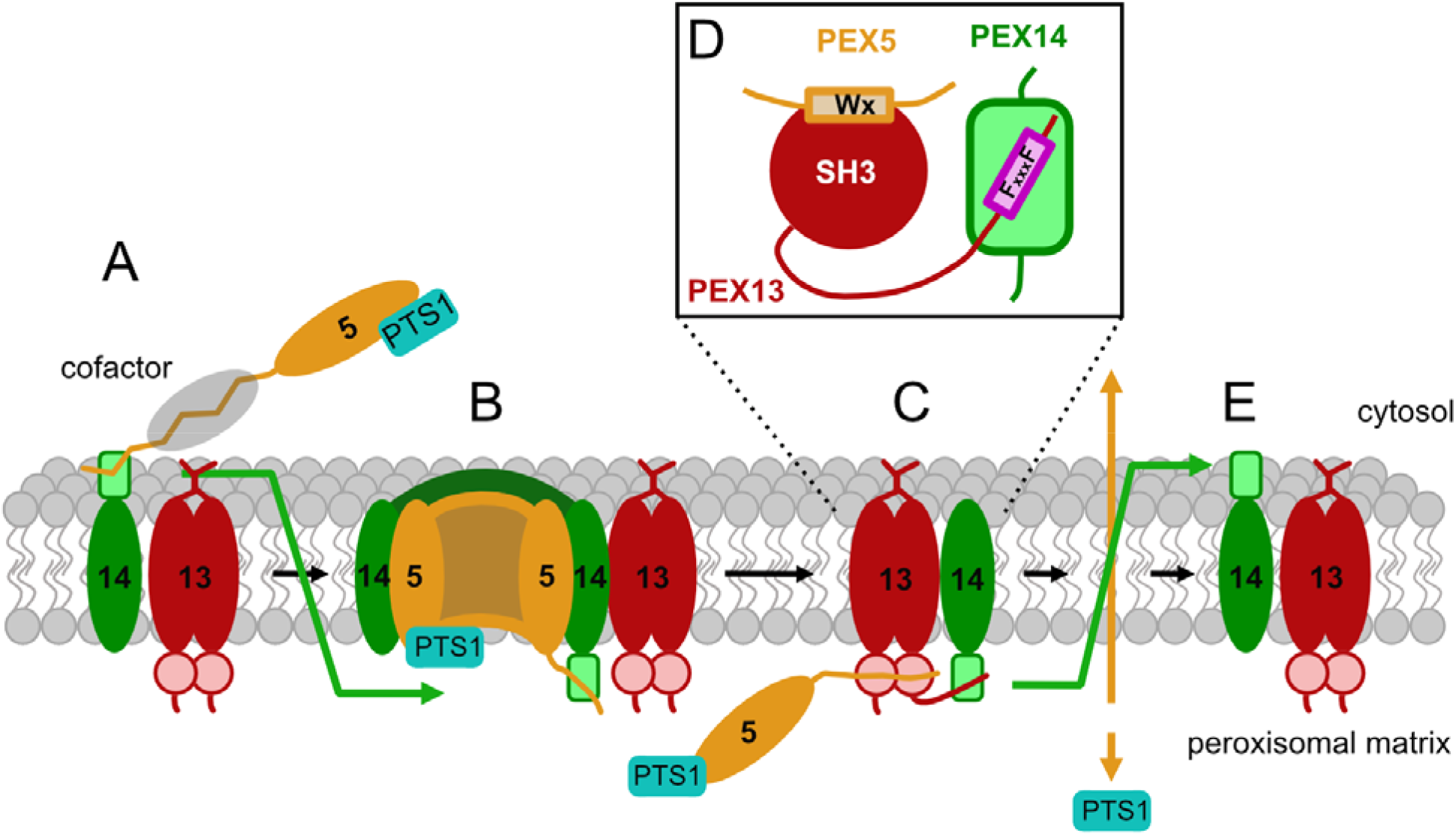
Proposed model for the role of PEX13 in peroxisomal matrix import. The import cycle can be explained with the following states. (A) High affinity binding of the cargo-loaded soluble receptor PEX5 to the cytosol faced N-terminal domain triggers release of potential PEX5 cofactors. (B) Pore formation of PEX5 with PEX14 and/or PEX13 mediates translocation of the cargo-bound PEX5 TPR domain and the PEX5 associated PEX14 NTD. (C) A PEX13 homo-oligomer replaces the PEX5/PEX14 interaction by avidity effects handing PEX5 over to PEX13 to form a transient complex. (D) Schematic representation of the interactions between PEX5, PEX14 and PEX13 in that state. (E) The transient complex dissembles when PEX5 is exported by the retrotranslocation channel (PEX2, PEX10 and PEX12). PEX14 flips orientation simultaneously with the recycling of PEX5.

In conclusion, our results highlight the importance of the PEX13 SH3 domain for peroxisomal import and demonstrate a regulatory function of the newly identified FxxxF motif via autoinhibitory interactions with the SH3 domain. The interaction network and competitive interactions of (di)aromatic peptide motifs with varying affinities in solution is modulated by differential localization and concentration of PEX5, PEX13 and PEX14 at the membrane and the presence of cargo bound to the PEX5 TPR domain. Future studies should analyze the PEX interaction network and complexes at the membrane to probe the model proposed here.

## Materials and Methods

### Molecular cloning

For recombinant expression in bacteria, the full length genes of human PEX13 (UniProtKB no. Q92968), human PEX14 (UniProtKB no. O75381) and human PEX5 (UniProtKB no. P50542) were optimized according to the codon usage of E. coli and synthesized by IDT (IDT Europe GmbH, Germany). These sequences were used as templates to generate PEX13 fragments PEX13 SH3 (261-346), FxxxF (261-383), SH3-CTR (261-403), 2GSc-FxxxF (chimera), GSc-W4 (chimera) as well as the PEX14 fragments PEX14 NTD (1-104), NTD_long_ (1-113) and PEX5 W0 (1-76) in a His_6_-SUMO-tag modified pETM13 vector (pETM13S). The cloning was done using site directed ligase independent mutagenesis (SLIM) (Chiu, 2004) in an extended version to implement inserts using the same fashion of short and tail primers. The vector backbone and inserts were amplified by polymerase chain reaction (PCR) amplification using the according short and tail primers (**Supplementary Table 2, 3**) to generate overlaps with sticky ends. The backbone amplificates and inserts were mixed with a 5-fold molar excess of insert and annealed during the SLIM cycle (Chiu, 2004). The annealed vector was directly transformed into DH10b cells for DNA amplification.

The same protocol was used to create the PEX13 constructs FL (1-403), trunc1 (1-264), trunc2 (1-346), trunc3, FxxxF to A5 and FxxxF to W4 substitution in the bi-cistronic mammalian expression vector pIRES2-EGFP-SKL. Primer are listed in **Supplementary table 4**.

### Protein sample preparation

PEX constructs were transformed into *Escherichia coli* BL21 (DE3) cells and expressed in LB or isotope-enriched M9 minimal medium. Uniformly ^15^N or ^15^N, ^13^C labeled proteins were expressed in H_2_O M9 minimal medium supplemented with 50 µg/ml kanamycin, 1g/liter [U-^15^N] ammoniumchloride and 2g/liter hydrated [U-^13^C] glucose as the sole sources of nitrogen and carbon, respectively. After transformation, single colonies were picked randomly and cultured in the medium of choice overnight at 37°C. On the next day, cultures were diluted to an optical density of 600 nm (OD_600_) of 0.1, and grown up to a OD_600_ of 0.4-0.6. Protein expression was induced with 0.5 mM IPTG and was carried out for 4h at 37°C.

The cells were harvested by centrifugation at 6000 g for 20 min at 4°C.For protein purification the cell pellets were resuspended in lysis buffer (50mM Tris pH 7.5, 300 mM NaCl, 20 mM imidazole) substituted with lysozyme (from chicken), DNAse and protease inhibitor mix (Serva, Heidelberg, Germany) and lysed by pulsed sonication (10 min, 40% power, large probe, Fisher Scientific model 550) followed by centrifugation at 38000 g for 45 min. All proteins were purified using gravity flow Ni-NTA (Qiagen, Monheim, Germany) affinity chromatography. The supernatant of the lysis was incubated with Ni-NTA beads (2ml/1l culture) for 20 min at 4 °C while rotating. Subsequently to incubation, the protein-laden beads were washed with 7 CV high salt buffer (50mM Tris pH 7.5, 750 mM NaCl, 20 mM imidazole) and 10 CV wash buffer (50 mM Tris pH 7.5, 300 mM NaCl, 20 mM imidazole). The elution was performed with 3-5 CV elution buffer (50 mM Tris pH 7.5, 300 mM NaCl, 500 mM imidazole). Dialysis and SUMO cleavage was executed over night at 4 °C in 20 mM Tris pH 7.5, 150 mM NaCl. Further purification was done with a reverse Ni-NTA column where the flow through containing the cleaved protein of interest was collected and concentrated for size exclusion chromatography using a Superdex S75, 16/600 (GE Healthcare, Rosenberg, Sweden). The size exclusion chromatography as last step of the purification was performed directly in NMR buffer.

### Peptide preparation

All 15-mer peptides used in this study were purchased from PSL (Peptide Specialty Laboratories GmbH, Heidelberg, Germany). Peptides delivered in TFA salt were dissolved in H2O and the pH adjusted to 7.5 using 1 M NaOH. If used for ITC, the peptides were dialyzed against ITC buffer (20 mM Tris pH 7.5, 50 mM NaCl).

PEX5 W1 ALSENWAQEFLAAGD

PEX5 W2 YNETDWSQEFISEVT

PEX5 W3 VSPARWAEEYLEQSE

PEX5 W4 TATDRWYDEYHPEED

PEX5 W5 AQAEQWAAEFIQQQG

PEX5 W6 GTSDAWVDQFTRPVN

PEX5 W7 AEAHPWLSDYDDLTS

### NMR spectroscopy

NMR data were collected on Bruker Avance III spectrometers operating at 500, 600, 800, 900 or 950 MHz, equipped with cryogenic probes. The sequential assignment of backbone resonances for PEX13 SH3-CTR was performed based on heteronuclear experiments such as ^1^H-^15^N-HSQC, HNCA, HN(CO)CA, CBCA(CO)NH, HNCACB, HNCO, HN(CA)CO, HN(CA)NNH and H(NCA)NN (Sattler M *et al*, 1999; Weisemann *et al*, 1993). {^1^H}-^15^N heteronuclear NOE (hetNOE) experiments (Farrow *et al*, 1994) were performed using the pulse sequence hsqcnoef3gpsi (Bruker, Avance version 12.01.11) with a 4.5 s interscan delay. NOE values are given simply by the ratio of the peak heights in the experiment with and without proton saturation (hetNOE = I_sat_/I_0_) (Renner *et al*, 2002). ^15^N HSQC-based *T*_1_ and *T*_2_ experiments used sequences developed from (Farrow *et al*., 1994) with water-control during the relaxation period in the *T*_1_ sequence using a cosine-modulated IBURP-2 pulse (Gairí *et al*, 2015) and modifications in the *T*_2_ sequences based on (Lakomek *et al*, 2012). For both *T*_1_ and *T*_2_ experiments 8 time points with delays of 80, 160, 240, 320, 400, 64, 800, 1000ms (*T*_*1*_) and 14.4, 28.8, 43.2, 57.6, 72.0, 86.4, 100.8, 115.2ms (*T*_2_) were measured respectively. NMR-Spectra were processed using Topspin (Bruker Biospin, Rheinstetten, Germany) or NMRPipe (Delaglio *et al*, 1995) and analyzed using CcpNMR Analysis 2.4.2 (Vranken *et al*, 2005).

All NMR experiments were performed at 298°K. PEX13 and PEX14 spectra were recorded in 20 mM Tris pH 7.5, 50 mM NaCl and 50 mM NaP pH 6.5 and 100 mM NaCl respectively.

For all titration experiments a reference protein concentration 100 µM was used. For protein/protein titrations, every titration point was prepared as individual sample to avoid dilution effects. Protein/peptide titrations such as titration of PEX5 (di)aromatic peptide motifs with high concentrated peptides (10-15mM) were performed in a single NMR tube. Ligands were added with increasing concentrations up to an excess of 8 fold. The chemical shift perturbation (Δδ_*avg*_) was calculated by using formula 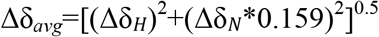. Chemical shift perturbations in **Fig. 5B** are illustrated with the sum of 10 residues which are affected upon (di)aromatic peptide motif binding. NMR chemical shift assignments are available at the BMRB, accession code: 51336.

### Isothermal Titration Calorimetry (ITC)

Isothermal titration calorimetry (ITC) measurements were performed as triplicates at 25°C using a MicroCal PEAQ-ITC (Malvern Instruments Ltd. U.K) calorimeter. Buffer conditions were 20 mM Tris pH 7.5, 50 mM NaCl. For all titrations a titrant dilution control experiment was performed and subtracted before the data were fitted to a one-site binding model using the Malvern Analysis software.

PEX13 SH3 at a concentration of 35-48 µM was titrated with purchased 15mer peptides containing diaromatic penta peptides from PEX5 or PEX5 1-76 (W0) at concentration of 0.7-0.6 mM and PEX13 350-403 (FxxxF) at a concentration of 0.9-1 mM. PEX14 (1-104) at a concentration of 20-60 µM was titrated with PEX5 W peptides or PEX5 W0 at concentration of 0.4 to 0.6 mM, PEX13 FxxxF at a concentration of 0.46mM and with PEX13 SH3-CTR at a concentration of 1 mM. The concentration of PEX14 was corrected with the fit, since it cannot be accurate measured at 280 nm owing to the extinction coefficient of only 1490.

ITC data are summarized in Tables 1, 2 and 3.

### X-ray crystallography

All crystals were grown using the vapor diffusion sitting drop method in 96 well plates. Therefore, the proteins were purified in 5 mM Tris pH 7.5, 50 mM NaCl and later screened using commercial crystallization sets (200 nl drops, 1:1 ratio). All proteins were crystallized at a concentration of 20mg/ml. The PEX 13 SH3 domain alone was crystallized in 0.01 M Zinc chloride, 0.1 M sodium acetate pH 5 and 20 % (w/v) PEG6000. PEX13 SH3 2GSc FxxxF chimera was crystallized in 0.2 M sodium chloride, 0.1 M Bis-Tris pH 6.5 and 25% PEG3350. PEX13 SH3 GSc PEX5 W4 chimera was crystallized in 0.2 M sodium sulfate, 0.1 M Bis Tris propane pH 6.5 and 20% (w/v) PEG3350. All crystals were cryoprotected using 25% ethylene glycol prior to flash freezing. Data of apo PEX13 SH3 and PEX13 SH3-2GCs-FxxxF chimera were collected at SLS beamline X06DA (Paul-Scherer-Institute, Villigen, Switzerland) while data of PEX13 SH3-GSc-W4 chimera was collected at beamline P11 at PETRA III (EMBL, Hamburg, DE).

Collected data were processed using CCP4i2 software suite (Potterton *et al*, 2018; Winn *et al*, 2011). XDS (Kabsch, 2010) was used for data indexing and reduction, aimless for data scaling, MOLREP (Vagin & Teplyakov, 2010) and PHASER (McCoy *et al*, 2007) for molecular replacement, COOT (Emsley & Cowtan, 2004) for model building and REFMAC5 (Murshudov *et al*, 2011) for refinement. Refined structures were uploaded to wwPDBdeposition using pdb_extract (Yang *et al*, 2004).

### Size exclusion chromatography - static light scattering (SEC-SLS)

SLS on PEX13 SH3-CTR was done using an OmniSEC Resolve and Reveal device (Malvern Panalytics. Malvern, Uk) equipped with a Superdex 75 increase 10/300 GL column (Cytiva). First 70 µl 2 mg/ml BSA standard (column calibration) and then 70 µl of 2 mg/ml PEX13-SH3-CTR in NMR/ITC buffer (20 mM Tris pH 7.5, 50 mM NaCl) was passed though the column with a constant flow of 0.3 ml/min. The concentration was monitored via absorption at 280nm and the refractive index. The molecular weight was calculated with the RALS signal using Omnisec software (version 11.01, Malvern Panalytics, Malvern, Uk).

### Computational Modelling

The structure of PI3K SH3 domain in complex with a PxxP ligand (PDB ID: 3I5R) was identified as similar to PEX13 SH3 domain by sequence search. Both domains were aligned and the PxxP ligand was copied to the PEX13 SH3 structure and subsequently mutated to PEX13 PxxP (TRVPPPIL) using Maestro (Schrodinger suite). The energy of the complex was then minimized using OPLS2005 force field and letting all residues in a radius of 4 Å freely rotate

### Multiple sequence alignments

#### Multiple sequence alignment of PEX13 SH3 with other human SH3 domains

First, a RCSB PDB databank search for structures of human SH3 domains was performed. Eight human SH3 domains from different proteins as well as yeast Pex13p SH3 domain were selected (**Supplementary Table 4**). Then the according sequences including the ± 10 flanking amino acids were selected and with the sequence from the human PEX13 SH3 aligned using clustalΩ (https://toolkit.tuebingen.mpg.de/tools/clustalo) and visualized using Jalview (version 2.11.2.0)

#### Multiple sequence alignment of PEX13 with mammalian sequences

Mammalian sequences which are similar to PEX13 were found using PSI-BLAST from ncbi blastp (https://blast.ncbi.nlm.nih.gov/) selecting the non-redundant protein sequences database with the organism restricted to mammals. The maximal target sequences were adjusted to 250 and the run started with preselected standards. From the hits, 186 sequences with a query coverage of minimum 90% were selected and aligned using clustalΩ (https://toolkit.tuebingen.mpg.de/tools/clustalo) before being visualized using ConSurf web server (Ashkenazy *et al*, 2016; Ashkenazy *et al*, 2010; Celniker *et al*, 2013) (https://consurf.tau.ac.il/).

### Cell culture

T-REx™293 (Invitrogen, USA) and T-REx293 PEX13KO cells (Ott et al., *in press*, https://doi.org/10.1515/hsz-2022-0223) were grown in Dulbecco’s Modified Eagle’s Medium high glucose (DMEM) supplemented with 10 % fetal calf serum, 4 mM L-glutamine, 100,000 U/l penicillin and 100 mg/l streptomycin at 37°C and 8.5% CO_2_. For cell transfections X-tremeGENE HP DNA Transfection Reagent (Roche, Germany) was used according to the manufacturer’s instructions.

### Fluorescence microscopy

To perform immunofluorescence microscopy cells were seeded on coverslips to appropriate density and fixed with 3% formaldehyde/D’PBS for 20 min. After membrane permeabilization with 1% Triton X-100/D’PBS for 5 min cells were incubated for 30 min in the primary antibody aPMP70 (rabbit, 1:500, Invitrogen) in D’PBS supplemented with 1% BSA. The incubation with the secondary antibody goat arabbit IgG (H+L) Alexa Fluor 594 (Invitrogen) was done for 10 min under light protection. Cells were mounted on glass slides with Mowiol 4-88 (Calbiochem, USA) supplemented with DAPI. Imaging was performed using the Axioplan 2 (Zeiss).

The quantification of import-competent cells was performed manually by cell counting in randomly taken images.

Quantification of rescued import was done over three biological replicates in total numbers. However, the import efficiency of PEX13 FL with 81% was normalized to 100% and import efficiencies from PEX13 variants were normalized to FL.

### Co-immunoprecipitation

To study interactions between PEX5 and PEX13 dynabeads were coupled with a mouse PEX5 antibody (Cizmowski et al., 2011), using the dynabeads™ Antibody Coupling Kit (Invitrogen, USA) according to the manufacturer’s instructions.

Cells were seeded on 10 cm dishes and transfected with different PEX13 truncations or mutations. 48 h after transfection cells were incubated in IP lysis buffer (25 mM Tris/HCl pH 7.4, 150 mM NaCl, 1 % NP-40, 5 % glycerol, cOmplete™ EDTA-free Protease Inhibitor Cocktail (Roche), 25 µg/ml DNase) for 15 min on ice. After removal of cell debris by centrifugation (13,000 rpm, 5 min, 4°C) equal amounts of lysates were incubated with dynabeads on a rotating disk for 1 h at 4°C. Next the dynabeads were washed three times with lysis buffer and bound proteins were eluted with 0.1 M Glycin pH 2.8. Samples were collected and analyzed by SDS-PAGE and immunoblotting using the following antibodies: rabbit aPEX5 (1:5000, rabbit aPEX13 (1:1000, Proteintech, Germany) (Fodor *et al*, 2015), chicken aPEX14 (1:1000, Ruhr-University Bochum), mouse aGAPDH (1:7500, Proteintech, Germany). Band intensities on immunoblots were quantified using densitometry (ImageJ, NIH).

## Data availability

- NMR chemical shift assignment data PEX13 SH3-CTR: BMRB 51336 (https://bmrb.io/data_library/summary/index.php?bmrbId=51336)
- Crystallographic data
  - PEX13 SH3: PDB (7Z0I) ()
  - PEX13 SH3-FxxxF chimera: PDB (7Z0J) ()
  - PEX13 SH3-PEX5-W4 chimera: PDB (7Z0K) ()

## 1 Acknowledgments

This work was supported by the Deutsche Forschungsgemeinschaft (FOR1905, project number 219314758, SA823/11-2, TP05) to M.S. and R.E. (ER178/6-2, ER178/7-2), and by the German Federal Ministry of Research and Education (BMBF) program “Targetvalidierung für die pharmazeutische Wirkstoffentwicklung” (GFTARV38) and BMBF VIP+ (03VP05531). F.D. acknowledges support by an EMBO Long-term Fellowship (ALTF 243**-**2018).

We thank Sam Asami, Mark Bostock, Gerd Gemmecker for support with NMR experiments. We acknowledge access to NMR measurements at the Bavarian NMR center. We are grateful to the X-ray Crystallography Platform at Helmholtz Zentrum München for support. We acknowledge the Paul Scherrer Institut, Villigen, Switzerland for provision of synchrotron radiation beamtime at beamline X06DA of the SLS and would like to thank Florian Dworkowski for assistance. Other synchrotron data were collected at beamline P11 operated by EMBL Hamburg at the PETRA III storage ring (DESY, Hamburg, Germany). We would like to thank Johanna Hakanpää for the assistance in using the beamline.

## 3 Author Contributions

MS, SG and RE designed the study. SG performed protein expression, biophysical and NMR experiments. SG and KZ performed X-ray crystallography, SG, KZ and GP analyzed crystallographic data. JO and WS planned and performed pulldown and complementation assays. SG and MS wrote the manuscript, all authors commented and approved the manuscript.

## Supplementary Material

**Supplementary Figure 1.**
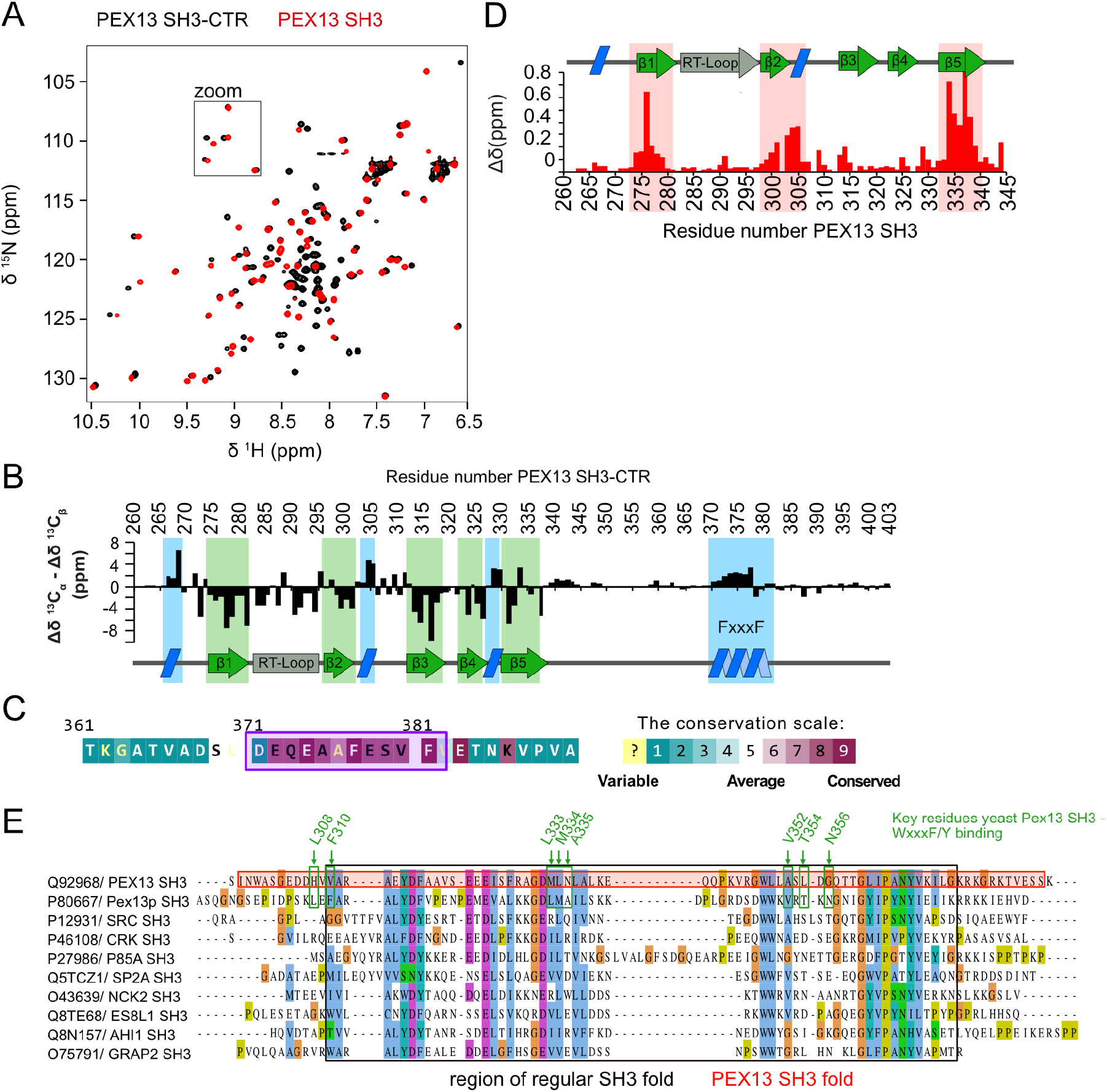
Conformation and NMR analysis of human PEX13 C-terminus. (A) Overlaid 2D spectra of PEX13 SH3-CTR (black) and PEX13 SH3 (red). A zoomed-view is shown in **Fig. 1** (B) ^13^C secondary chemical shifts (Δδ^13^Cα - Δδ^13^Cβ) of PEX13 SH3-CTR (261-403) support the typical β-sandwich fold of the SH3 domain and the presence of a short α-helical motif comprising the FxxxF motif. (C) Sequence alignment with 186 mammalian PEX13 sequences plotted with ConSurf web server (Ashkenazy *et al*, 2016; Ashkenazy *et al*, 2010; Celniker *et al*, 2013) shows high conservation of the FxxxF motif (purple box). (D) chemical shift perturbations plotted on the PEX13 SH3 sequence with secondary structure indicated on top. (E) Sequence alignment of human PEX13 SH3 with yeast PEX13 SH3 and SH3 domains from human SRC, CRK, AHI1, NCK2, ES8L1, GRAP2, P85A, and SPD2A (with known structures). The sequence alignment was done with the ± 10 residues flanking the SH3 fold. The black box indicates the regular SH3 fold and the red box the complete fold of human PEX13 SH3. Green arrows indicate key residues for yeast Pex13 SH3 / WxxxF/Y binding. Note that those residues are poorly conserved from yeast to human.

**Supplementary Figure 2.**
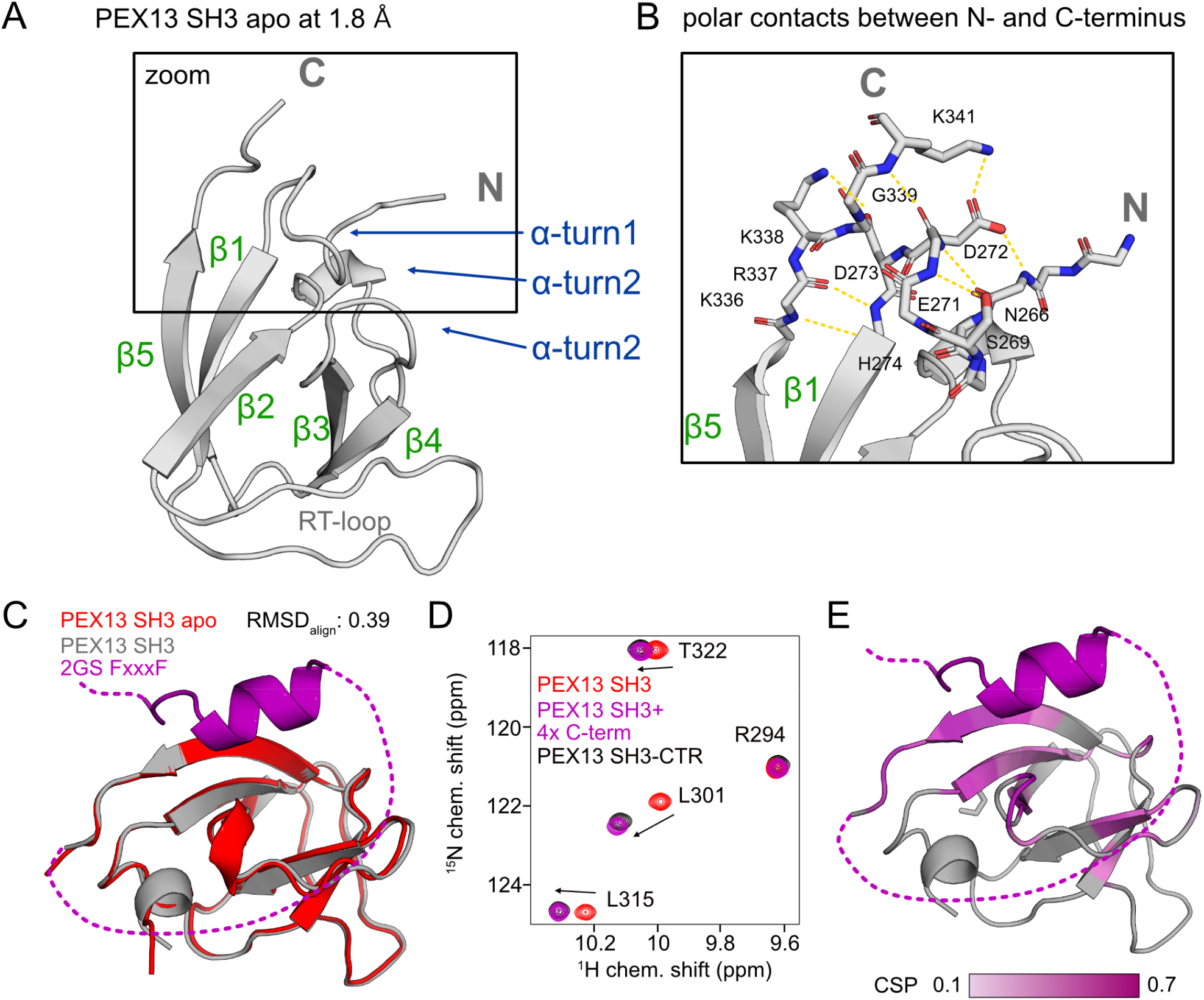
Structural features of apo PEX13 SH3 and confirmation of PEX13 SH3 – FxxxF structure in solution. (A) Apo structure of PEX13 SH3 solved with 1.8 Å resolution. (B) Zoomed-view showing a network of hydrogen bonds in the on the N – and C-terminal regions (yellow). (C) Superimposition of the structures PEX13 SH3 (red) and PEX13 SH3 (gray) in complex with FxxxF motif (purple). (D) Overlay of ^1^H, ^15^N correlation spectra of PEX13 SH3 (red) PEX13 SH3-CTR (black) and PEX13 SH3 titrated with FxxxF peptide (350-403) (purple). PEX13 SH3 titrated with FxxxF peptide represents the native spectrum of PEX13 SH3-CTR. (E) Chemical shift perturbations from PEX13 SH3 / FxxxF titration mapped on the PEX13 SH3-FxxxF complex structure.

**Supplementary Figure 3.**
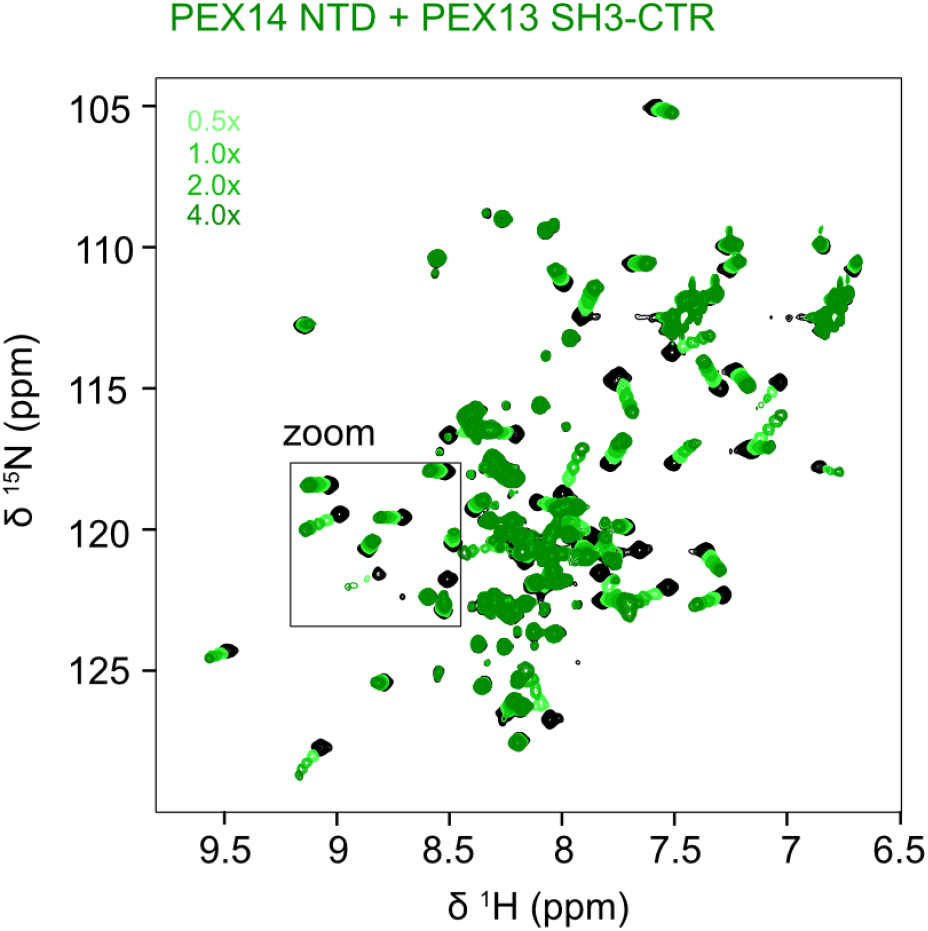
Titration of 15N PEX14 NTD with PEX13 SH3-CTR. (A) Overlaid2D spectra of PEX14 NTD (black) titrated with increasing concentrations PEX13 SH3-CTR (261-403) (green scale). Zoom is shown in Fig. 4.

**Supplementary Figure 4.**
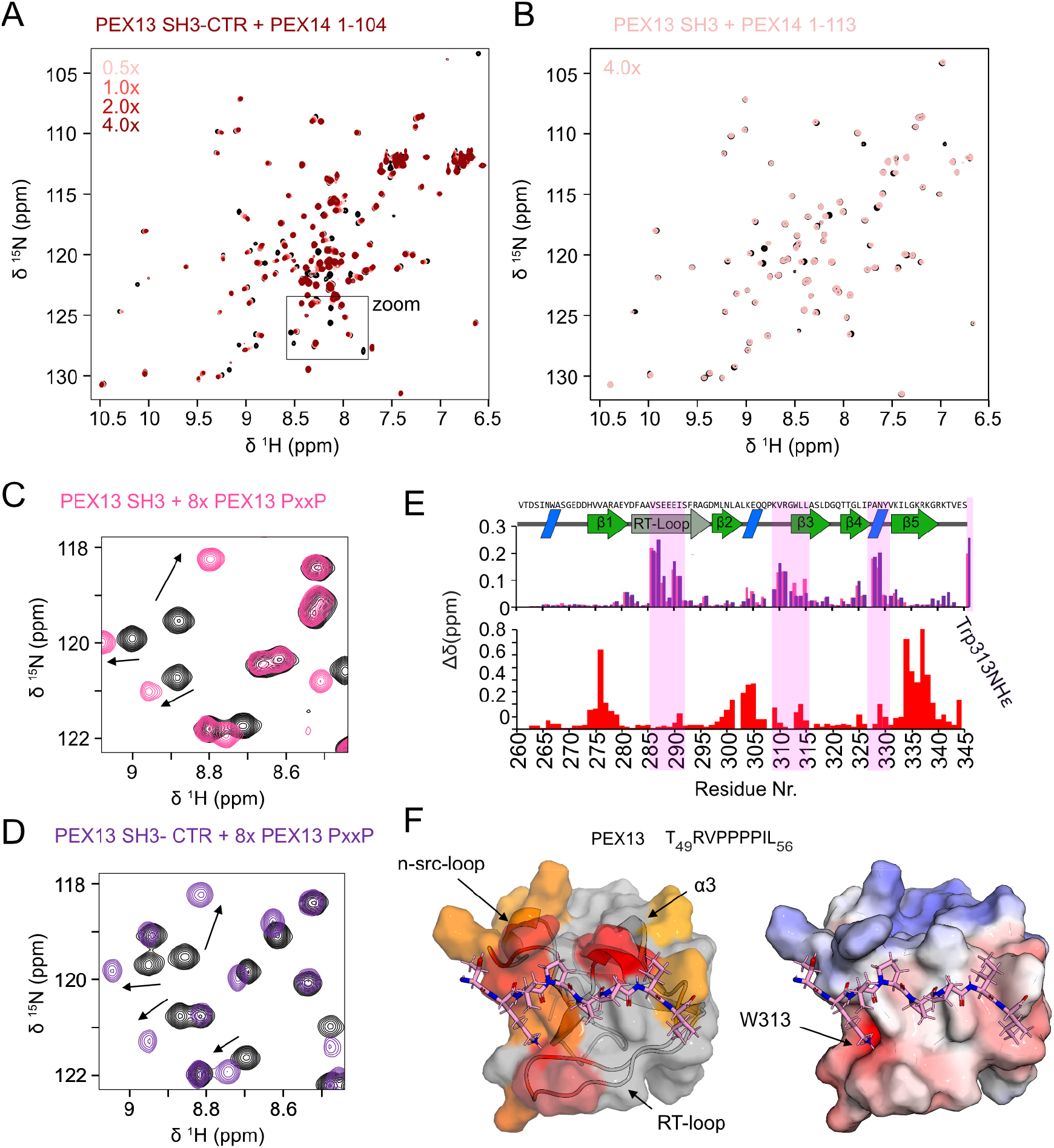
Interactions of PEX13 SH3 with PxxP peptides. (A) Overlaid 2D spectra of PEX13 SH3-CTR (black) titrated with increasing concentrations PEX14 NTD (1-104) (red scale). Zoom is shown in main figure. (B) 2D spectra of PEX13 SH3 with 4x excess of PEX14 NTDlong (1-113) (C) 2D spectra of PEX13 SH3 (black) with 8x excess of PEX13 PxxP peptide (49-61) (pink). (D) 2D spectra of PEX13 SH3-CTR (black) with 8x excess of PEX13 PxxP peptide (49-61) (dark purple) (E) Chemical shift perturbations from PxxP titration on SH3 (pink) and SH3-CTR (dark purple) (upper panel) and from FxxxF titration on SH3 (red/bottom panel) mapped on the PEX13 SH3 sequence demonstrating non-overlapping binding sites (F) Structure of PEX13 SH3 with modelled PEX13 PxxP (49-56) represented with transparent surface showing chemical shift perturbations (yellow to red gradient) from NMR titrations (left panel) and electrostatic surface (right panel). PxxP sequence shown above. Binding of PEX13 PxxP (49-61) induces typical CSPs in the regions of RT-loop, n-src-loop and α3 (α10) (Shi *et al*, 2012).

**Supplementary Figure 5.**
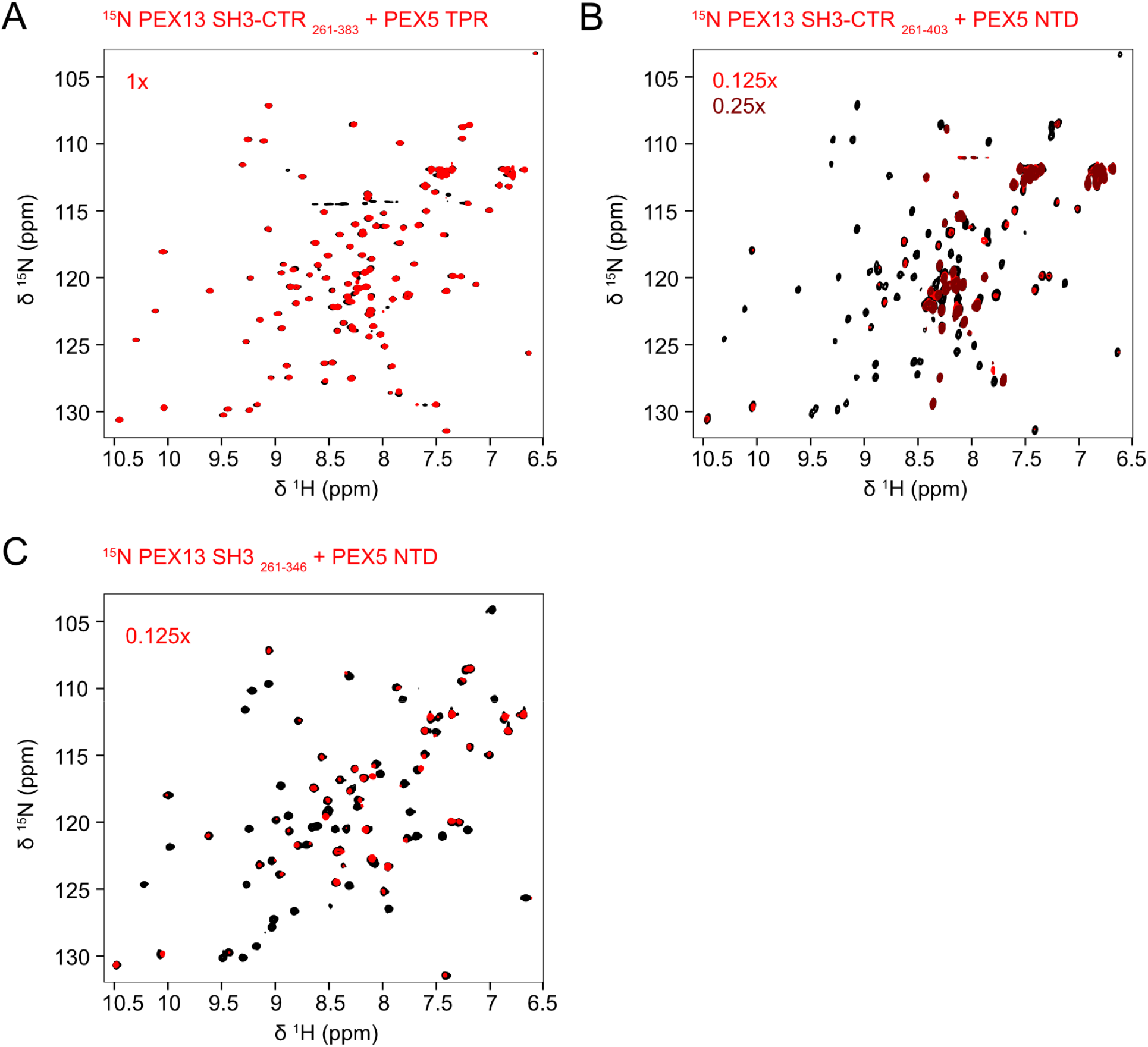
Titration of ^15^N PEX13 SH3-CTR or PEX13 SH3 with PEX5 TPR, PEX5 NTD or PEX13 C-terminal peptide. (A) Overlaid 2D spectra of PEX13 SH3-CTR (261-383; missing the last 20 amino acids; black) titrated with equimolar concentration of PEX5 TPR domain (red). (B) Overlaid 2D spectra of PEX13 SH3-CTR (black) titrated with 0.125x (red) or 0.25x (dark red) excess of PEX5 NTD. Notably, resonances experience excessive line-broadening at 0.125 (1/8) ligand concentration. (C) Overlaid 2D spectra of PEX13 SH3 (black) titrated with 0.125x (red) excess of PEX5 NTD. The SH3 titration experiment shows very similar line-broadening effect as seen with PEX13 SH3-CTR indicating binding of PEX5 NTD to PEX13 SH3.

**Supplementary Figure 6.**
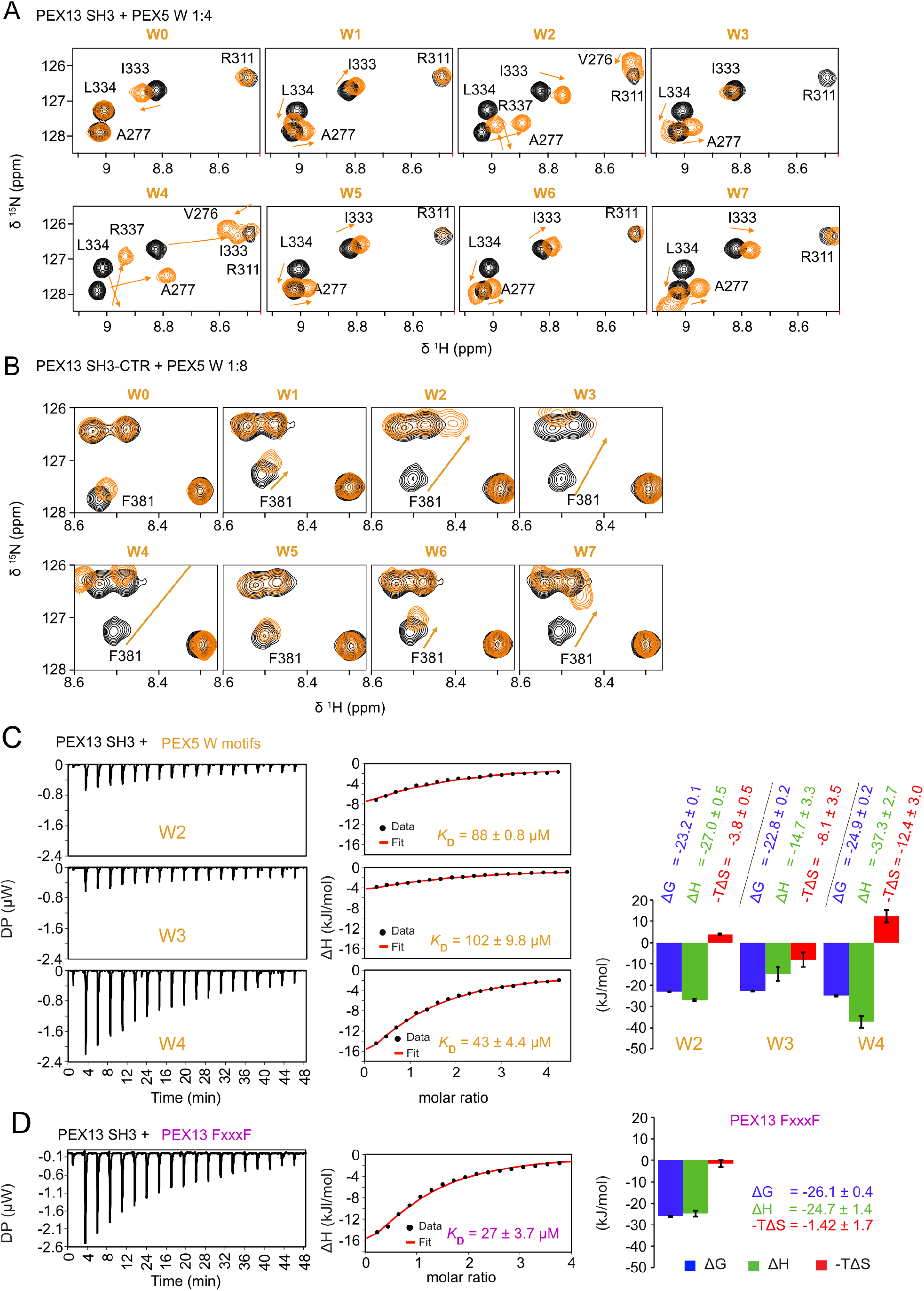
(Competitive) binding of PEX13 SH3 and SH3-CTR with PEX5 W peptides. (A) Overlaid 2D spectra of PEX13 SH3 (black) and with 4x excess of PEX5 W peptides (orange). (B) Overlaid 2D spectra of PEX13 SH3-CTR (black) and with 8x excess of PEX5 W peptides (orange). (C) ITC titration of PEX13 SH3 with PEX5 W2, W3 and W4 (D) ITC titration of PEX13 SH3 with FxxxF peptide (350-403)

**Supplementary Figure 7.**
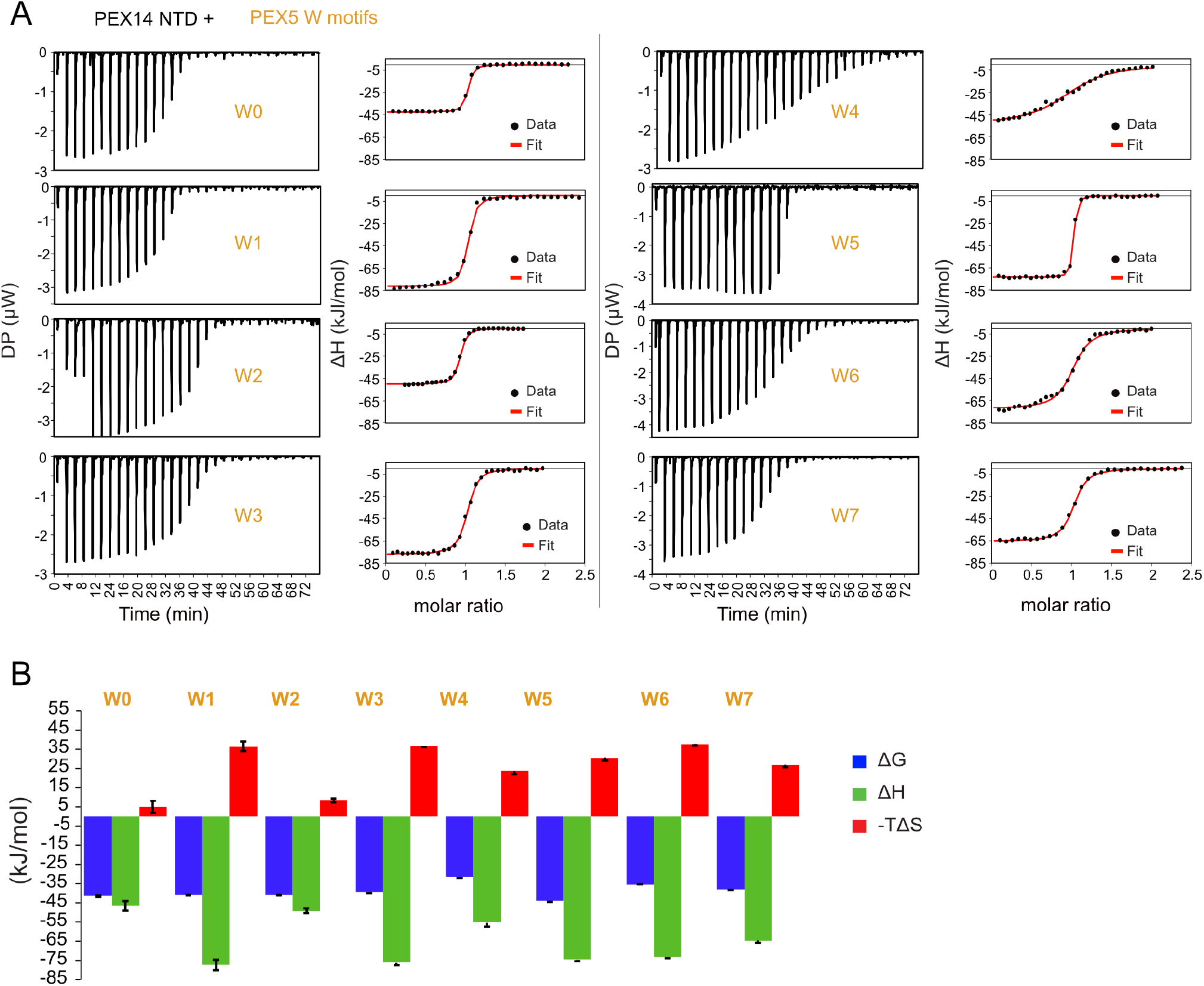
Isothermal titration calorimetry of PEX14 NTD/ PEX5 W interactions. (A) ITC titration of PEX14 NTD (1-104) with PEX5 W (-like) motifs. (B) Energetic contribution of ITC titration PEX14 NTD (1-104) with PEX5 W (-like) motifs.

**Supplementary Figure 8.**
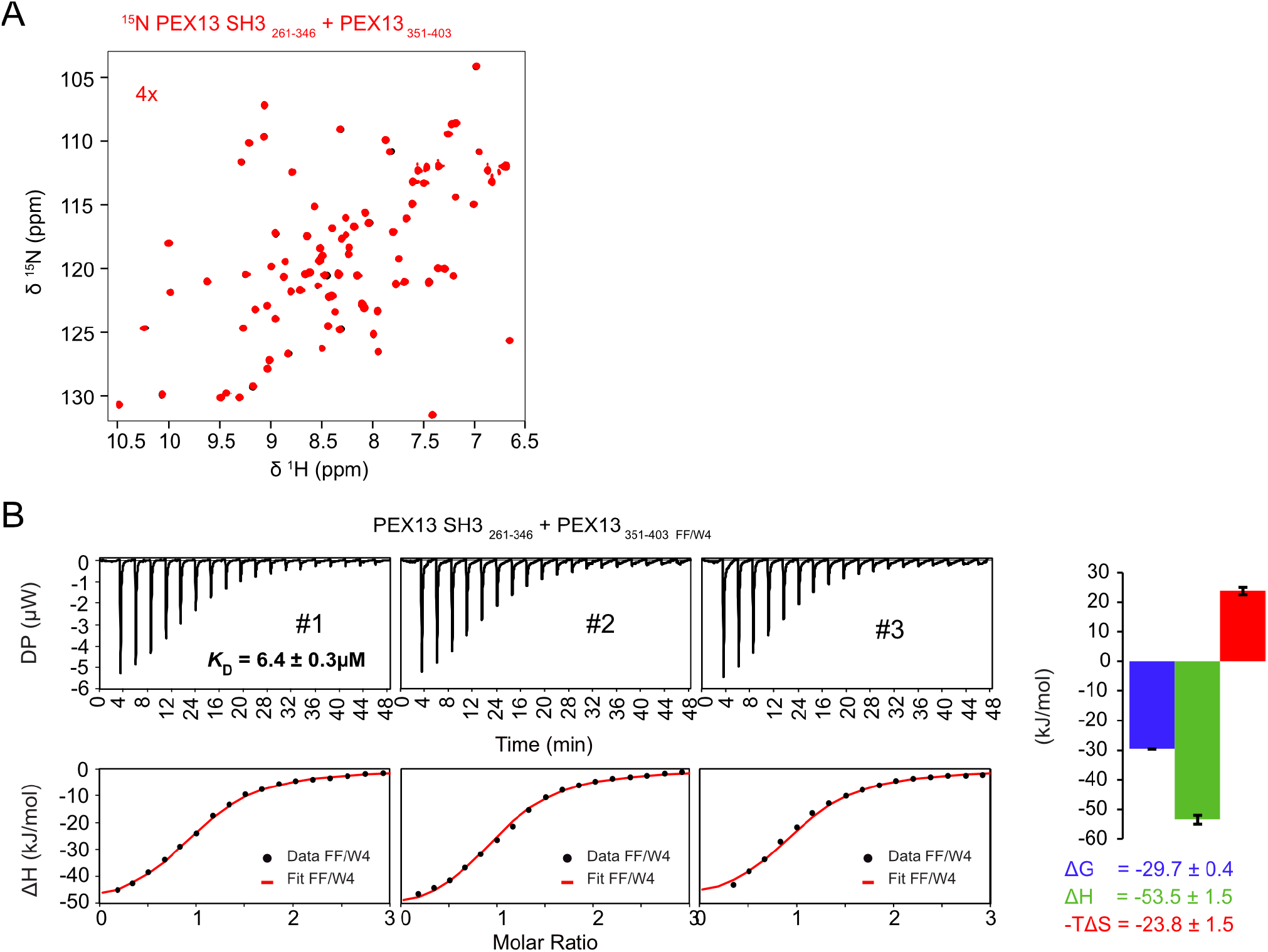
Binding of PEX13 SH3 domain to PEX13 C-term A5 or W4 core motif mutations. (A) Overlaid 2D spectra of PEX13 SH3 (black) titrated with 4x (red) excess of PEX13 C-terminal peptide (351-403) with the FxxxF motif mutated to AAAAA indicate no binding activity. (B) ITC experiments with the SH3 domain and the C-terminal peptide FxxxF to PEX5 W4 show a clear binding event with a dissociation constant *K*D ∼ 6 µM. The curve was fitted to a one binding site model with N=1.

**Supplementary Figure 9.**
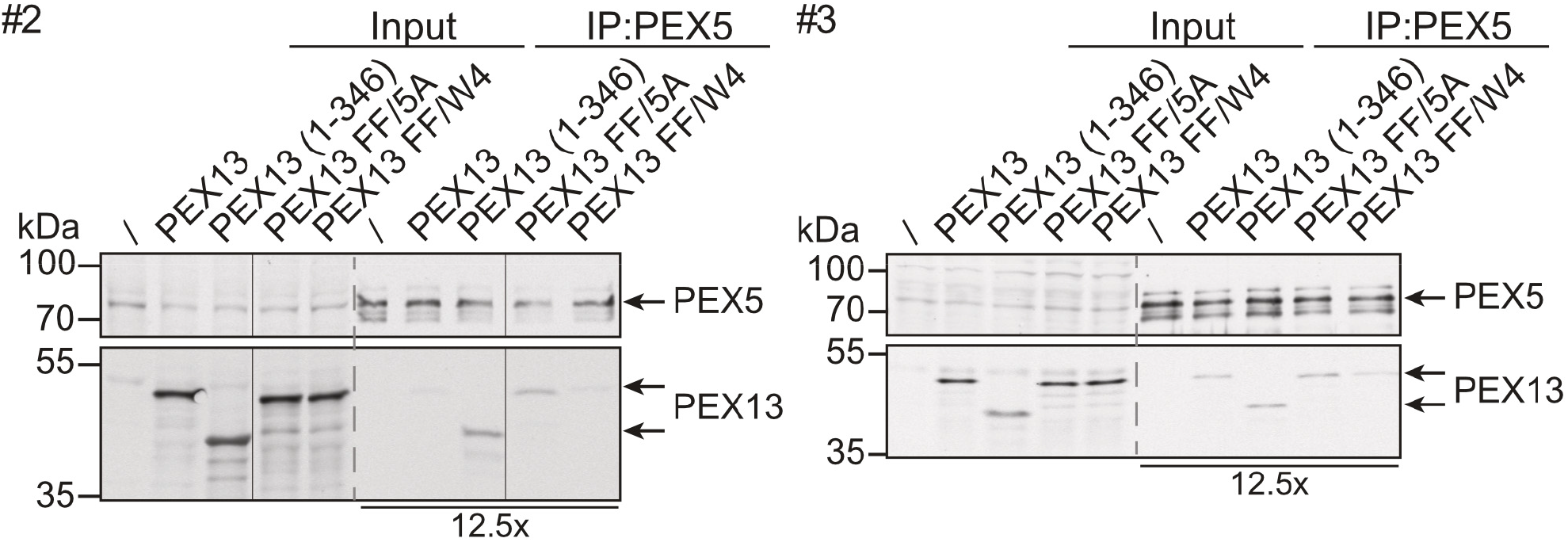
Repetitions two (#2) and three (#3) of the PEX13 pull-down experiments. PEX13 FL, PEX13 1-346 and the two mutations FF/A5, FF/W4 (B) were expressed in T-Rex PEX13 KO cells. The cell lysates were subjected pull down procedures and affinity purification with PEX5 antibody and analyzed via SDS PAGE and immunoblotting.

## Supplementary Tables

**Supplementary Table 1.** Crystallographic data for PEX13 SH3, PEX13 SH3 2GS FxxxF and PEX13 SH3 W4 structures.

|  | PEX13 SH3 | PEX13 SH3 2GS<br>FxxxF | PEX13 SH3 GS W4 |
| --- | --- | --- | --- |
| PDB entry | 7Z0I | 7Z0J | 7Z0K |
| Space group | I 41 | P 31 2 1 | P 31 2 1 |
| Cell parameters (Å, °) | 43.94 43.94 86.78<br>90.00 90.00 90.00 | 86.80 86.80 65.43<br>90.00 90.00 120.00 | 87.66 87.66 66.11<br>90.00 90.00 120.00 |
| **Data collection** |  |  |  |
| Wavelength (Å) | 0.999995 | 1.000029 | 1.0332 |
| Temperature (K) | 100 | 100 | 100 |
| Resolution range (Å) | 43.39-1.80<br>(1.84-1.80) | 49.36-2.3<br>(2.38-2.3) | 49.86-2.3<br>(2.38-2.3) |
| Total no. of observed reflections | 104090 (6202) | 260870 (25799) | 468378 (25836) |
| Number of unique reflections | 7660 (462) | 12998 (1250) | 13362 (1275) |
| R<sub>merge</sub> | 0.082 (1.443) | 0.282 (2.426) | 0.495 (10.645) |
| R<sub>pim</sub> | 0.034 (0.588) | 0.091 (0.774) | 0.118 (3.458) |
| CC1/2 | 1 (0.738) | 0.997 (0.652) | 0.992 (0.588) |
| I/σ(I) | 19.7 (2.1) | 11.8 (1.6) | 17.0 (1.8) |
| Completeness (%) | 99.9 (99.5) | 100 (100) | 100 (100) |
| Multiplicity | 13.6 (13.4) | 20.1 (20.6) | 35.1 (20.3) |
| Wilson B-factor (Å<sup>2</sup>) | 26.2 | 35.67 | 45.11 |
| **Refinement** |  |  |  |
| Rwork (%) | 0.182 | 0.197 | 0.193 |
| Rfree (%) | 0.229 | 0.224 | 0.217 |
| No. Of atoms | 619 | 1326 | 1359 |
| Ligand atoms | 11 | 12 | 0 |
| Solvent atoms | 51 | 75 | 61 |
| **Model quality** |  |  |  |
| RMS (Bonds) | 0.0087 | 0.0133 | 0.0156 |
| RMS (angles) | 1.498 | 1.9 | 2.05 |
| Ramachandran favored (%) | 96.05 | 97.56 | 96.93 |
| Ramachandran allowed (%) | 3.95 | 2.44 | 3.07 |
| Ramachandran outliers | 0 | 0 | 0 |
| Rotamer outliers (%) | 6.35 | 6.62 | 10 |
| Clashscore | 0.8 | 2.63 | 3.71 |

**Supplementary Table 2.**
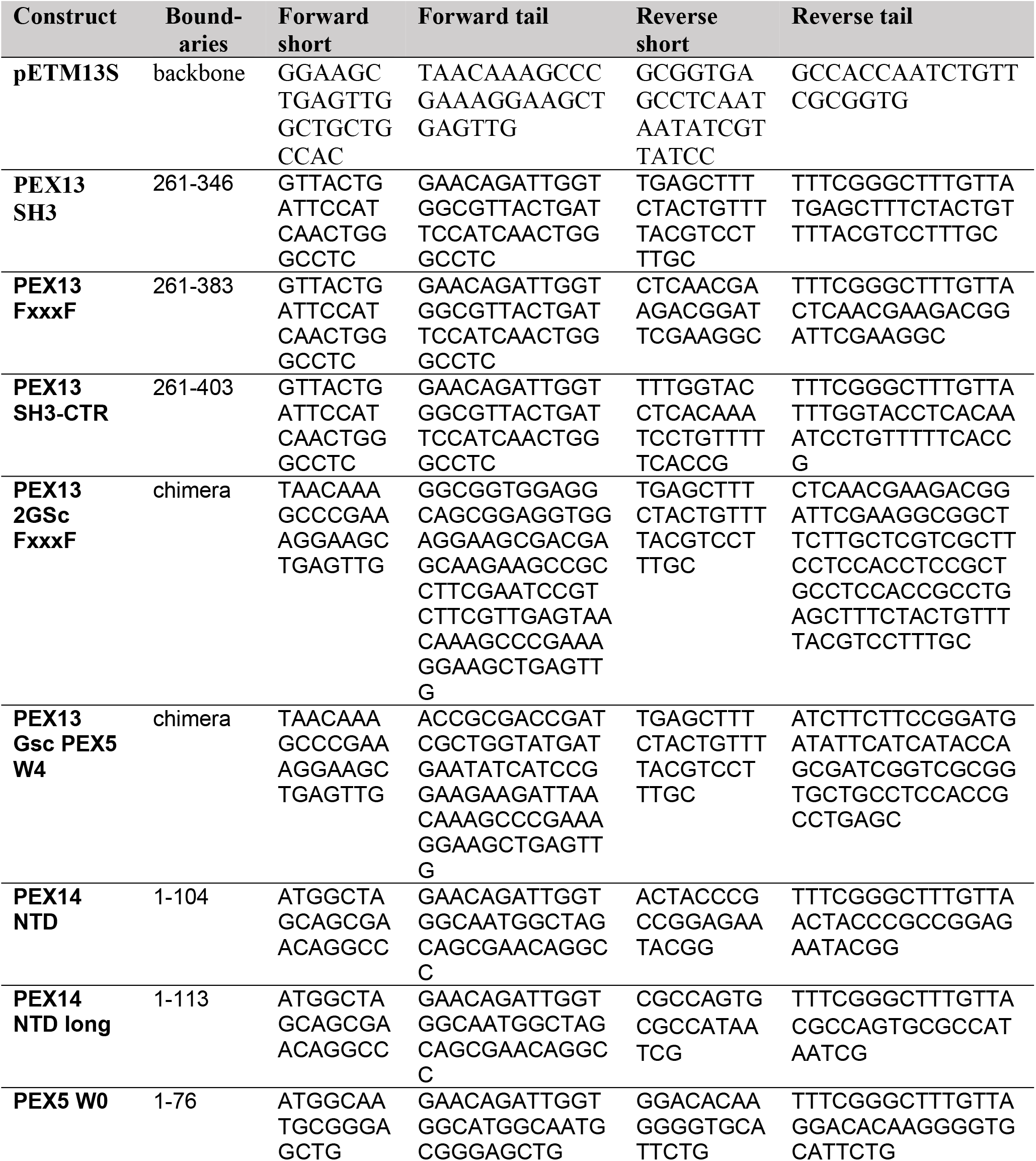
Primer list for cloning into pETM13S vector.

**Supplementary Table 3.**
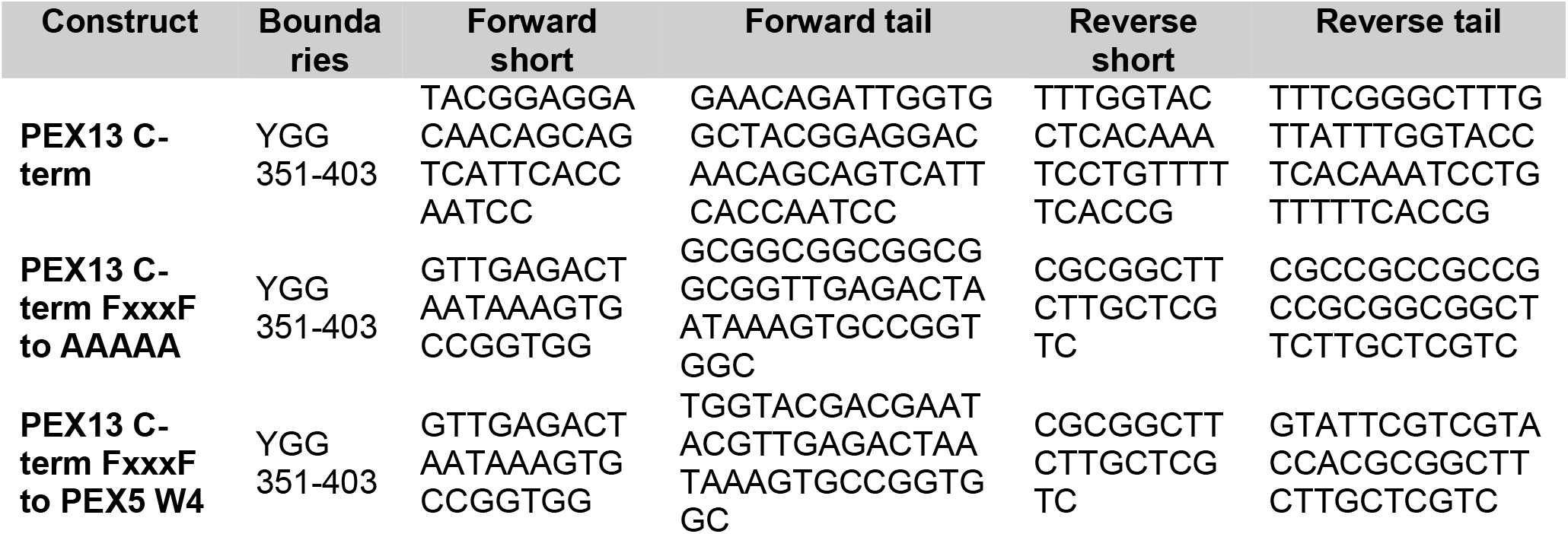
Primer list for cloning PEX13 C-term variants into pETM13S vector.

**Supplementary Table 4.**
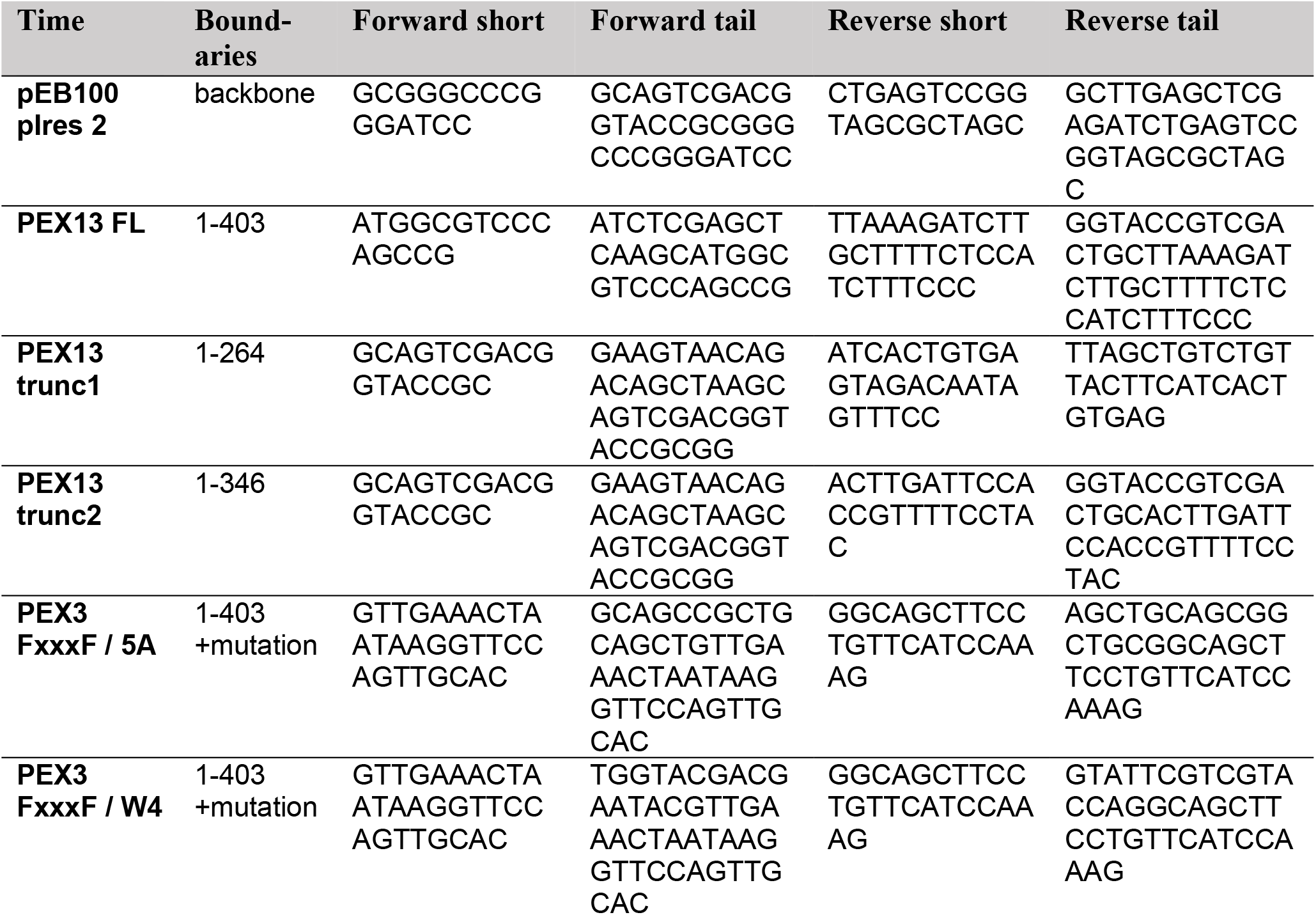
Primer list for cloning into bi-cistronic vector pIRES2 - GFP.

**Supplementary Table 4.**
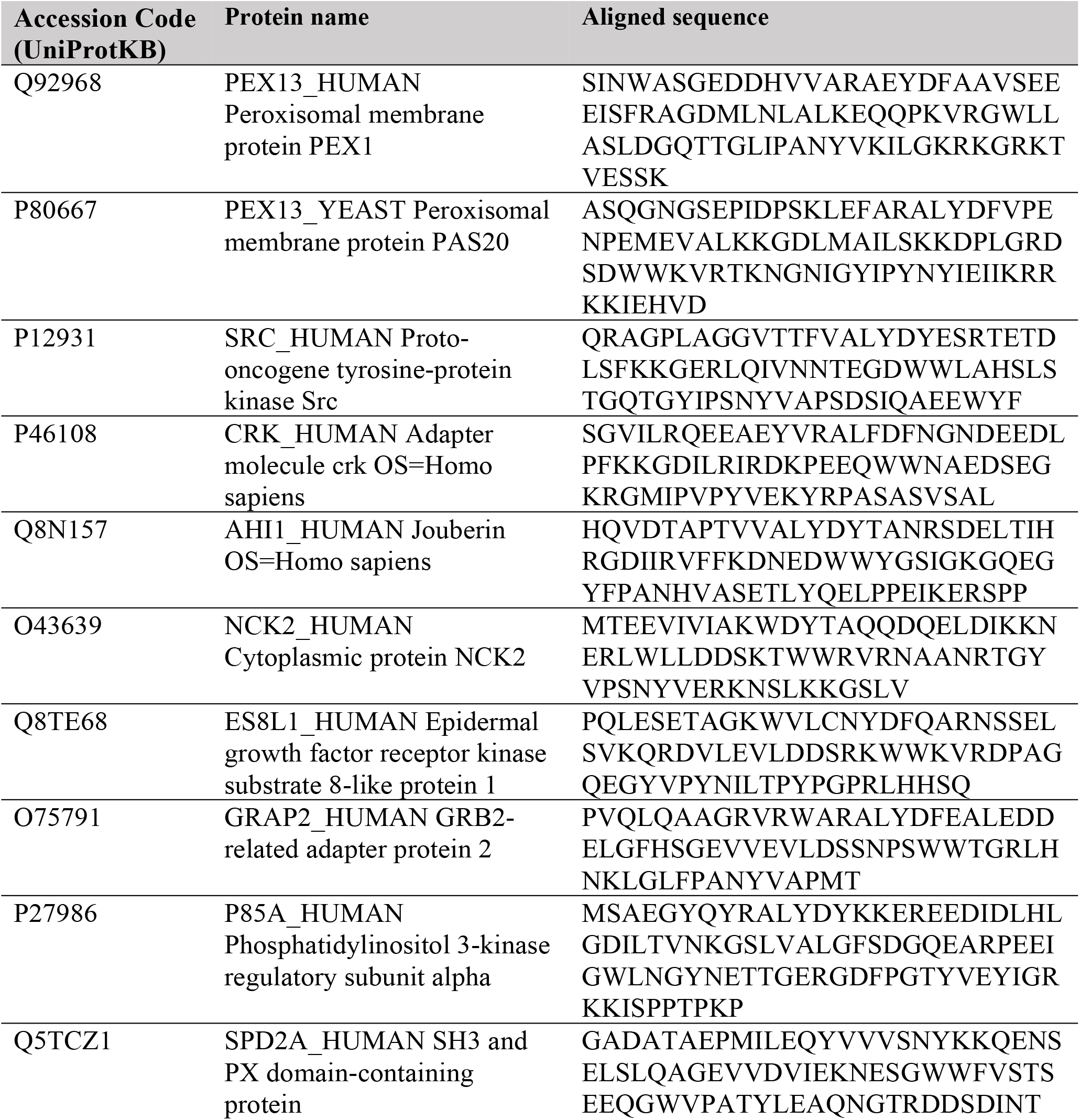
Sequences of human SH3 domains for MSA.

